# Blood flow simulation and uncertainty quantification in extensive microvascular networks: Application to brain cortical networks

**DOI:** 10.1101/2024.12.05.627123

**Authors:** Peter Mondrup Rasmussen

**Affiliations:** Department of Clinical Medicine, Center of Functionally Integrative Neuroscience, Aarhus University, Aarhus, Denmark

## Abstract

0.1

Spatially resolved simulation models of microcirculatory blood flow facilitate a detailed understanding of microcirculatory phenomena at the micrometer scale by capturing heterogeneity in blood flow. These models combine physical laws, empirical descriptions of the blood’s complex rheological behavior, and in-vivo/ex-vivo imaging of the microvasculature. However, imaged areas often only partially represent self-contained tissue regions, leading to numerous vessels crossing boundaries and strongly influencing simulated blood flows through imposed boundary conditions. Selecting appropriate boundary conditions is challenging due to the heterogeneity of pressures and blood flows, resulting in significant uncertainties.

This study addresses two key methodological aspects of spatially resolved blood flow simulations: selecting appropriate boundary conditions and quantifying the impact of boundary condition uncertainties on simulated hemodynamic variables. An adaptive method for assigning appropriate pressure boundary conditions is proposed and rigorously evaluated in extensive brain cortical networks against reference data from an established blood flow simulation model. A probabilistic approach is adopted to assess the impact of boundary condition uncertainties on blood flow simulations. The adaptive method is further integrated into a Bayesian calibration framework, inferring distributions over thousands of unknown pressure boundary conditions and providing uncertainty estimates for blood flow simulations.

The adaptive method, which is straightforward to implement and scales well with extensive microvascular networks, produces hemodynamic simulations consistent with reference data, yielding depth-dependent pressure profiles and layer-wise capillary blood flow profiles consistent with previous studies. These phenomena are demonstrated to generalize also to biphasic blood flow simulation models incorporating in-vivo viscosity formulations. The uncertainty analysis further reveals a novel spatially heterogeneous and depth-dependent pattern in blood flow uncertainty. It is anticipated that the adaptive method for pressure boundary conditions will be useful in future applications of both forward and inverse blood flow modeling, and that uncertainty quantification will be valuable in complementing hemodynamic predictions with associated uncertainties.

**0.2 Author summary:** This research focuses on improving the accuracy of blood flow simulations in tiny blood vessels, known as microvascular networks. These simulations help understand how blood moves through the smallest vessels in the body, crucial for studying various health conditions. However, accurately simulating blood flow is challenging because imaged areas often don’t capture entire tissue regions, leading to uncertainties.

I developed an adaptive method for setting boundary conditions in these simulations. Due to its adaptive nature, the method can be applied to microvascular networks from various types of tissue, making it broadly applicable. This method was tested extensively using data from brain cortical networks and produced reliable results, proving its validity and scalability to extensive networks.

Additionally, probabilistic approaches were used to assess how uncertainties in boundary conditions affect the simulations. A key contribution is the integration of the adaptive method into a Bayesian calibration framework. This framework assimilates simulations with observations and infers distributions over thousands of unknown boundary conditions, providing uncertainty estimates for blood flow simulations.

The proposed adaptive method and uncertainty analysis are expected to be valuable for future studies of microvascular blood flow, improving both the accuracy of the simulations and the understanding of the associated uncertainties.

## 1. Introduction

Microvascular networks are complex structures composed of numerous interconnected tiny blood vessels[1]. These networks support normal microcirculatory function by bringing blood into close proximity with tissue, thereby facilitating the transport of oxygen and nutrients into tissue while simultaneously removing waste products. Microcirculatory variables are inherently heterogeneous, both structurally and functionally, including strong variations in red blood cell (RBC) velocities and blood oxygenation[2–4]. These heterogeneities are inevitable consequences of inherent network characteristics, including vessel segment connectivity, variations in segment lengths and diameters, and rheological phenomena governing microscale blood flow[2, 5]. This microscale heterogeneity, in turn, significantly influences biophysical phenomena at macroscale, such as oxygen and nutrient delivery to tissue and clearance of waste products[4, 6–9].

Despite their complexity, microvascular networks are well-organized and tightly regulated to ensure an adequate distribution of blood across tissue[1, 10–16]. However, disruption in the microcirculation can lead to altered oxygen and nutrient delivery to tissue and clearance of waste products causing cellular malfunction and various diseases[16, 17]. For instance, impaired microcirculatory mechanisms have been implicated in neurodegenerative diseases[16, 18–21], highlighting the need to understand microcirculatory phenomena at network level to uncover underlying pathophysiological changes[4, 17].

Investigating microcirculatory phenomena is challenging due to its small spatial scale. Despite this, a broad range of experimental measurement techniques have been developed, allowing for in-vivo measurement of key physiological parameters such as blood flow velocities and oxygenation levels in blood and in tissue[10, 22]. These in-vivo measurements offer valuable quantitative data about normal microcirculatory function and facilitate the evaluation of changes in various diseases. However, current in-vivo measurement techniques have limitations. For instance, these techniques are often restricted to a sparse set of spatial locations, which can result in an incomplete characterization of the tissue region being studied. Consequently, extrapolation from average values for “typical vessels” to network properties can lead to substantial errors due to the presence of strong heterogeneities and correlations between microcirculatory hemodynamic parameters[3]. Additionally, there is a series of important salient but physiologically significant parameters that cannot be assessed by current measurement techniques[1, 3].

Biophysical modeling is a powerful approach that complements in-vivo studies, providing a deeper understanding of microcirculatory function[23, 24]. For example, biophysical modeling enables in-silico experiments where parameters affecting microcirculatory function can be carefully controlled, allowing for the assessment of their impact on microcirculatory phenomena[24, 25]. Furthermore, assimilating biophysical modeling with in-vivo measurements into a comprehensive analysis facilitates a network-oriented and model-based interpretation of measurements and provides insights into latent microcirculatory parameters that are otherwise unobservable[3, 26, 27].

Spatially resolved microcirculatory modeling aims to provide a comprehensive biophysical understanding of microcirculatory phenomena at the micrometer scale[23]. By incorporating spatial resolution, these models can capture the heterogeneity in microcirculatory variables, providing an accurate representation of microcirculatory blood flow and solute transport phenomena[3]. Furthermore, the spatial resolution in these models corresponds closely to the measurement scale of in-vivo microscopic measurements. As a result, the predictions made by these models can aid in hypothesis development and provide a foundation for planning future in-vivo experiments. These experiments can then, in turn, contribute to the ongoing model development and enhancement in a cyclical process.

Spatially resolved microcirculatory blood flow models are usually constructed using a combination of a set of physical laws, empirical descriptions of the blood’s complex rheological behavior, and in-vivo/ex-vivo imaging of the microvasculature[1, 28–33]. Microvasculature often forms a continuous network within tissue. However, the imaged area often only partially represents a self-contained tissue region, leading to numerous vessels crossing the boundaries of the imaged area[34]. It is widely recognized that simulated blood flows are heavily influenced by imposed boundary conditions at these cut vessels, making the selection of appropriate boundary conditions a persistent challenge that requires considerable attention[26, 28–31, 33–38]. A significant contributing factor to this challenge is the heterogeneity of pressures and blood flows throughout the microvasculature[2, 3], leading to substantial uncertainties regarding the boundary conditions[26].

Recently, we introduced a Bayesian probabilistic approach for assimilating spatially resolved blood flow simulation models with hemodynamic observations, accounting for errors in the observed data as well as uncertainties in model parameters, including boundary conditions and parameters of rheological descriptions[26, 39]. Boundary pressures were treated as uncertain parameters, and by calibrating model predictions against in-vivo measurements or literature-derived data, the Bayesian approach offered an effective means for inferring distributions over the unknown boundary conditions while simultaneously providing a formal means for quantitatively assessing the impact of these uncertainties on model predictions.

The present study continues along this line of research and addresses the complexities in modeling blood flows in extensive microvascular networks, with thousands of boundary conditions, and in quantifying the impact of the inevitable boundary condition uncertainty on model predictions through the following methodology: (i) An adaptive method for setting appropriate pressure boundary conditions is proposed. (ii) The predictions made by this new method are subjected to rigorous quantitative validation in extensive rodent cortical microvascular networks, against reference model predictions based on an established blood flow simulation model[25, 31]. (iii) A probabilistic approach is adopted to assess the influence of uncertainty in pressure boundary conditions on blood flow predictions in forward simulations based on reference pressure boundary conditions and in simulations obtained with the proposed method. (iv) The new adaptive method for setting pressure boundary conditions is incorporated into our Bayesian calibration framework[26, 39], and the utility of the framework is evaluated by implementing it with three different viscosity models[40–42]. Similarity in model predictions of hemodynamic variables and in uncertainty estimates among blood flow simulation models incorporating these three viscosity models as well as their similarity with the reference model predictions is evaluated. (v) Depth-dependent pressure profiles and laminar capillary blood flow profiles are examined across models, and the depth-resolved analysis is further complemented by an assessment of blood flow uncertainty along the underlying blood flow paths.

## 2. Materials and methods

This paper reports the author’s own work and ideas. Spell- and grammar-checking tools in Microsoft Word and Microsoft Copilot were employed for spell-checking, grammar-checking, and suggesting rephrasing of smaller pieces of original text to improve readability. All corresponding text was subsequently reviewed and edited by the author. No original content was generated by these tools, and the author takes full responsibility for the content of the publication. This declaration is provided to maintain transparency and integrity in accordance with publishers’ and institutional guidelines.

### 2.1. Overview of material and methods

The material and methods section presents the governing equations underlying the blood flow simulation model and outlines the method used for modeling non-uniform hematocrit. The adaptive method for setting pressure boundary conditions is described, and the process of integrating this method into the blood flow simulation model is also described. Following this, a probabilistic approach that enables a quantitative evaluation of the impact of boundary condition uncertainty on blood flow simulations is described followed by a description of our Bayesian calibration framework. Lastly, the microvascular networks and numerical experiments used in the quantitative model evaluation are described.

Throughout the text, matrices will be denoted with capital boldface letters ***A***, vectors by small boldface letters ***a***, scalars by small letters *a*, while *A*_*i*,*j*_ will denote the element in the i’th row and j’th column in ***A***.

### 2.2. Blood flow modeling

#### 2.2.1. Equations governing blood flow

Blood flow simulations are based on mass conservation, information on network topology, vessel segment morphology, and empirical descriptions of blood’s complex rheological behavior. Let a microvascular network comprise *n* nodes and *m* segments. Segment blood flow rate can be assumed to be governed by Poiseuille’s Law[28, 33] yielding the following relationship between segment blood flow rate ***q*** and node pressures ***p***

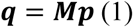

***q*** is an *m* × 1 vector of segment blood flow rates, ***p*** is an *n* × 1 vector of node pressures, and ***M*** is a sparse *m* × *n* matrix of segment hydraulic conductance (inverse resistance) with non-zero elements 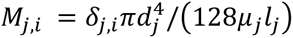, where δ_*j*,*i*_ is +1 (resp. -1) if node *i* is start node (resp. end node) of segment *j*, whereas *d*_*j*_, *l*_*j*_, and μ_*j*_ denote segment diameter, length, and effective viscosity, respectively. A segment’s blood flow rate *q*_*i*_ is positive (resp. negative) if the flow direction is from the segment’s start node to its end node (resp. end node to its start node). The effective viscosity and hence the hydraulic resistance depends non-linearly on both diameter and hematocrit and can be described by empirical models[1]. Mathematical details of three empirical models of effective viscosity[40–42] used in the present study are available in Section 1 in S1 File.

Blood flows at interior nodes sum to zero (mass conservation), which together with imposed pressure or flow boundary conditions encoded in ***b*** leads to[35]

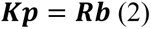

In Eq 2, ***K*** = ***LM*** + ***J*** + ***N*** is an *n* × *n* sparse matrix. ***L*** is an *n* × *m* sparse matrix with non-zero elements *M*_*i*,*j*_ = 1 (resp. *M*_*i*,*j*_ = −1) if node *i* is an interior node and is the start node (resp. end node) of segment *j*. ***J*** is an *n* × *n* sparse matrix with non-zero diagonal elements *J*_*i*,*i*_ = 1 if a pressure boundary condition is imposed at boundary node *i*. ***N*** is an *n* × *n* zero matrix and ***R*** is an *n* × *n* identity matrix. At this point, the two matrices ***N*** and ***R*** have no influence in Eq 2 and could be removed without loss of generality. However, their coefficients will be modified in the proposed adaptive method for pressure boundary conditions, and the matrices are hence introduced here to maintain notational consistency. ***b*** is an *m* × 1 sparse vector, with non-zero elements *b*_*i*_ = −|*q*_*i*_ | (resp. *b*_*i*_ = |*q*_*i*_ |) if blood inflow rate (resp. outflow rate) |*q*_*i*_ | is imposed at boundary node *i*, or *b*_*i*_ = *p*_*i*_ if boundary pressure *p*_*i*_ is imposed at boundary node *i*. After imposing pressure or flow boundary conditions at boundary nodes, the system of equations in Eq 2 can be solved efficiently by numerical solvers designed for sparse linear systems[43]. After solving Eq 2 for node pressures, the resulting node pressures can be substituted into Eq 1 to yield segment blood flows.

#### 2.2.2. Phase separation

In a microvascular network, vessel segments often form a highly connected network with numerous branchpoints[1, 12] of which bifurcations are the predominant type[1]. In downstream branches of a bifurcation, it is well-established that red blood cells (RBCs) are distributed in a non-uniform manner, the phenomenon known as the phase separation effect[1, 44]. The distribution of RBCs among the daughter branches is determined by several factors including the hematocrit of the mother branch, the distribution of blood flow between the daughter branches, and the vessel diameters. The specific phase separation model[30, 39, 41, 44] used in this study is detailed in Section 1 in S1 File. Phase separation results in extensive heterogeneity in hematocrit throughout microvascular networks[28–31]. This heterogeneity, in turn, affects the distribution of blood flow due to the influence of hematocrit on blood’s effective viscosity. To account for this interdependence of blood flow distributions and hematocrit distributions in blood flow simulations, an iterative procedure has been established[28, 30]. This procedure involves alternating between estimating the blood flow distribution based on a given hematocrit distribution and estimating the hematocrit distribution based on the resulting blood flow distribution. Algorithmic details of the iterative procedure used in this study can be found in Section 2 in S1 File.

### 2.3. An adaptive method for assigning pressure boundary conditions

#### 2.3.1. Model definition

An adaptive method for assigning pressure boundary conditions is proposed. The rationale behind the method is a reproducing property in the model, ensuring that statistical properties of the respective boundary conditions mirror those of a set of interior nodes. Specifically, the method establishes pressure boundary conditions in relation to some average pressure characteristics of a set of interior nodes. The method’s adaptive nature, which eliminates the need for a specific declaration of a reference pressure level, therefore allows it to inherently adapt to differences that may exist between different microvascular networks. Simultaneously, the heterogeneity in pressures, as observed in microvascular networks[3, 29–31, 33], can be included by imposing specific pressure deviations, at individual boundary nodes, relative to this reference level.

Mathematically, the pressure at boundary node *i* is defined relative to a weighed sum of pressures at a set of interior reference nodes

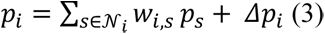

with 𝒩_*i*_ being a set of interior reference nodes each with pressure *p*_*s*_, *w*_*i*,*s*_ > 0 are normalized weighting coefficients, i.e. ∑_*s*∈𝒩*i*_ *w*_*i*,*s*_ = 1, and *Δp*_*i*_ is an imposed pressure deviation relative to the weighted sum of reference pressures.

There are many strategies for defining the set 𝒩_*i*_ of interior reference nodes for a given boundary node *i*. Reference nodes could for example be identified based on some measure of similarity, for example node type similarity (e.g., arteriolar, venular, or capillary node), morphological similarity, or geometrical similarity (e.g., similar cortical depth or similar topological location). Note that while different boundary nodes may have identical sets of interior reference nodes, the actual boundary pressures may still differ due to the application of pressure deviations *Δp*_*i*_ at the level of individual boundary nodes in Eq 3.

There are likewise many strategies for defining the weighting coefficients *w*. These could, for example all be equal, for a given boundary node, and the resulting sum over reference nodes in Eq 3 would then yield the average reference node pressure. Another example could be that the weighting coefficients are defined based on some distance measure such as Euclidean distances or the shortest path-distances between a given boundary node and the individual reference nodes.

After defining sets of interior reference nodes and corresponding weighting coefficients, the challenge of defining boundary pressures has now been re-cast into a challenge of defining the pressure deviations *Δp*_*i*_ relative to the set of reference nodes. These relative pressures could for example be sampled from some random distribution to mimic the pressure heterogeneities observed in microvascular networks, expert knowledge may be used to impose systematic deviations relative to the reference node pressures, or they could be inferred in data assimilation by using a model calibration technique[26]. In the following, the pressure deviations in Eq 3 will be referred to as *relative pressure boundary conditions* to distinguish these from conventional pressure boundary conditions.

#### 2.3.2. Incorporating the adaptive method into the governing system of equations

The adaptive method for defining pressure boundary conditions relative to a set of reference nodes, Eq 3, can be incorporated into the governing system of equations, Eq 2, by modifying the elements in the matrices ***J*** and ***N***. The matrix ***J*** will contain ones at diagonal places corresponding to boundary nodes governed by either conventional pressure boundary conditions or relative pressure boundary conditions. ***N*** will encode information about the sets of interior reference nodes and their associated weighting coefficients in Eq 3 as follows. For a given boundary node *i*, governed by a relative pressure boundary condition, the matrix ***N*** will have non-zero elements *N*_*i*,*s*_ = −*w*_*i*,*s*_ corresponding to its reference nodes *s* ∈ 𝒩_*i*_. Finally, the relative pressure deviation *Δp*_*i*_ will be encoded in ***b*** by *b*_*i*_ = *Δp*_*i*_. It is thereby straightforward to incorporate the adaptive method for pressure boundary conditions in Eq 3 into the governing system of equations, Eq 2, which can then be solved as usual.

#### 2.3.3. Preserving sparsity

The linear systems in Eqs 1 and 2 may experience a significant reduction in sparsity if numerous boundary nodes are subjected to relative pressure boundary conditions, and many reference nodes are utilized concurrently. This decrease in sparsity can subsequently result in a substantial increase in the computational effort needed to solve the system of equations, Eq 2. However, it is often reasonable to assume that multiple boundary nodes may have identical sets of reference nodes. Even under this assumption, the pressures at these boundary nodes may still vary due to the application of the relative pressure deviation *Δp*_*i*_ at the level of individual boundary nodes. In one extreme scenario, all boundary nodes governed by relative pressure boundary conditions could all have identical sets of reference nodes, which could be defined by all interior nodes with equal weight, for example. In this case, the pressures at these boundary nodes would be defined relative to the average interior node pressure. Alternatively, sets of reference nodes could be identical for boundary nodes that possess a common structural property. For instance, groups of boundary nodes could be defined based on a specific geometric property, such as similar cortical depth, which is the approach used in this study.

Consider 𝒢_*g*_ as a set of boundary nodes that all have identical sets of reference nodes, with the weighting coefficients for individual reference nodes also being identical across these boundary nodes. Sparsity can then be increased by designating one of the boundary nodes *i* ∈ 𝒢_*g*_ as a primary node and the remaining boundary nodes *i*^′^ ∈ 𝒢_*g*_, *i*′ ≠ *i* as secondary nodes. This requires modifying the definition of non-zero elements in ***N*** and the corresponding elements in ***R*** in Eq 2. For the specific group of boundary nodes 𝒢_*g*_, ***N*** will then contain non-zero elements *N*_*i*,*s*_ = −*w*_*i*,*s*_ only for the primary node *i*. Rows corresponding to secondary nodes will, on the other hand, only have one non-zero elements at *N*_*i*′,*i*_ = −1. The elements in ***R*** need to be adjusted accordingly and will, in addition to the ones along the diagonal, contain non-zero off-diagonal elements *R*_*i*′,*i*_ = −1 for secondary nodes. The enhanced sparsity of ***N*** has a crucial implication since it can lead to a tremendous reduction in the computational effort needed for solving the Eq 2. This reduced computational effort is particularly important when the system of equations must undergo multiple solutions, such as during hematocrit iterations or in calibration analysis.

### 2.4. Probabilistic uncertainty quantification

#### 2.4.1. Uncertainty quantification in simulation models

In the context of modeling, the blood flow simulation model can be regarded as a forward model[24, 26, 45]. When a particular set of model parameters, denoted by ***θ*** (for example, boundary conditions), is prescribed, the model’s states ***h***(***θ***) (for example, segment blood flows, velocities, pressures, and hematocrits) can be predicted. Boundary conditions are subject to an inevitable presence of uncertainty which, consequently, introduces uncertainties into model predictions[26]. By treating the uncertain parameters ***θ*** as random variables with associated probability distributions, this inherent uncertainty and the resulting uncertainty in model predictions can be effectively captured[46, 47]. By adopting rules of probability, the probabilistic approach thereby offers a formal and statistically rigorous means for quantifying model uncertainties.

For example, considering segment blood flows as the variable of interest (i.e. blood flows in ***q***(***θ***) constitutes the states ***h***(***θ***) above), the probabilistic approach enables assessment of how uncertainties in boundary conditions translate into uncertainties in these blood flows. Formally, the probability distribution that governs segment blood flows, given the uncertainty in boundary conditions, can be inferred. Uncertainties in boundary conditions lead to varying degrees of uncertainties in blood flows throughout a microvascular network due to segment connectivity and the heterogenous morphology across segments. Consequently, some segments’ blood flow may exhibit a substantial level of uncertainty, while others may display lesser uncertainty. This uncertainty is captured by the probability distribution governing segment blood flows which can be summarized by various summary statistics. The expectation

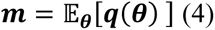

provides a measure of central tendency, representing the average or expected value of individual segments’ blood flow under uncertainty, aiding in understanding the typical behavior of the system. The covariance matrix

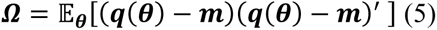

on the other hand, quantifies the spread or dispersion around this expected value, thereby capturing the inherent uncertainty in blood flows. Its off-diagonal elements convey information on between-segment covariance, while its diagonal elements provide insight into individual segments’ variance. Consequently, the square root of the *j*’th diagonal element in ***Ω*** corresponds to the standard deviation *σ*_*j*_ that governs the simulated blood flow in segment *j*. The mean and covariance thereby collectively offer a comprehensive statistical characterization of the system’s behavior under uncertainty, facilitating a robust and informed analysis of the blood flow simulations.

Additional metrics may further be derived. For instance, the coefficient of variance for individual segments’ blood flows can be calculated from the average blood flows in ***m***, Eq 4, and standard deviations derived from the diagonal elements in ***Ω***, Eq 5. A large standard deviation for a given segment, relative to its mean, indicates high uncertainty in blood flow, and blood flow may even exhibit opposite directions under this uncertainty. The following metric, termed the direction agreement rate (DAR), was established to capture this effect and is defined by

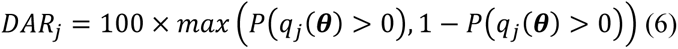

Here, *P*(*q*_*j*_ (***θ***) > 0) is the probability that the segment’s blood flow, which is a random variable, follows the direction from the segment’s start node to its end node. The DAR metric thereby quantifies the rate at which a segment’s blood flow direction agrees with its predominant direction. A large DAR, close to 100%, indicates that a segment’s blood flow is governed by a small dispersion, corresponding to a small standard deviation *σ*_*j*_ relative to the mean *m*_*j*_. Conversely, a small DAR, close to 50%, occurs with large dispersion. The DAR metric thus provides a straightforward, yet insightful and interpretable means for quantitively assessing the uncertainties in blood flow simulations.

The probabilistic approach to uncertainty quantification depends on methods to determine the distribution governing segment blood flows, considering uncertainty in boundary conditions. The following two sections will describe methods for inferring this distribution. First, a simplified approach will be applied to linear forward simulation models, followed by a more sophisticated method, anchored in Bayesian statistics, which takes observed data or literature-derived data into account.

#### 2.4.2. Uncertainty quantification in forward modeling

Often, specific flow or pressure boundary conditions are imposed in blood flow simulation models. These boundary conditions might, for instance, be derived from measured data[3], derived from literature data[3, 28, 30, 31], through domain-specific reasoning[28, 30], by numerical optimization and calibration techniques[26, 35, 37, 48], or by embedding the network within an extensive synthetic network[31]. Assuming specific boundary conditions have been defined, the objective of the following is then to establish a straightforward yet effective approach to quantify the extent to which uncertainties in these imposed boundary conditions lead to uncertainties in the simulated blood flows.

The governing equations Eqs 1 and 2 can be combined to yield

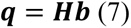

where ***H*** = ***MK***^**−1**^***R***. As previously described, incorporating the phase separation effect into the flow simulation model results in an interdependence between ***q*** and ***H***. However, under the simplifying assumption of a known hematocrit field, (for example, uniform hematocrit[27, 34, 49]), the mathematical formulation in Eq 7 emphasizes that segment blood flows are linear transformations of the pressure or blood flow boundary conditions in ***b***. This property will be utilized in the following approach for quantifying blood flow uncertainties related to boundary condition uncertainty.

Let the set of row indices corresponding to boundary nodes in ***b*** be partitioned into two sets 𝒜 and ℬ. These two sets consist of boundary nodes assumed to be associated with uncertainty and boundary conditions assumed to be known without associated uncertainty, respectively. Eq 7 can then be written as

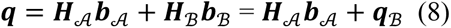

where ***H***_*x*_ and ***b***_*x*_ are composed of columns and rows in ***H*** and ***b***, respectively, corresponding to the indices contained in the two sets 𝒜 and ℬ. From a probabilistic perspective, assume that the uncertainty governing the boundary conditions at nodes in 𝒜 is modeled by a Gaussian distribution with mean ***b***_𝒜_ and covariance ***Σ***_𝒜_ i.e. 𝒩(***b***_𝒜_, ***Σ***_𝒜_). That is, the expected values of these boundary conditions are the same as defined in ***b*** whereas the additional boundary condition uncertainties are described in terms of the covariance matrix ***Σ***_𝒜_. It follows from statistical theory[46], that variables resulting from a linear transformation of Gaussian variables, Eq 8, are also governed by a Gaussian distribution, i.e.

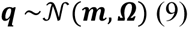

Segment blood flows are thereby random variables with the expected value

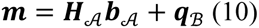

identical to the blood flows in Eqs 7 and 8. The additional uncertainties in blood flows, induced by the uncertainties in boundary condition, are described by the covariance matrix

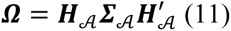

In a microvascular network, a certain degree of covariance between the boundary condition variables is expected due to the network’s structural connectivity. However, this covariance information is complex and not readily accessible. Instead, a pragmatic approach, which was employed in the present study, is to treat the covariance matrix ***Σ***_𝒜_ as a diagonal matrix. Hence, boundary condition uncertainties are described in terms of uncertainty related to individual boundary variables. Although this is a simplification, it’s important to note that the covariance matrix ***Ω***, which governs the distribution of segment blood flows in Eq 9, accumulates uncertainty contributions from all individual boundary variables by filtering these through the matrix ***H***_𝒜_ as seen in Eq 11. It should be noted that the probabilistic approach to uncertainty quantification, as outlined above, is related to classical sensitivity analysis, as further described in Appendix, Section 5.1.

The description above, Eqs 7-9, hinges on the assumptions of a linear simulation model, i.e. fixed hematocrit, and the application of a Gaussian distribution to characterize boundary condition uncertainty. These choices facilitate an analytical deviation of the distribution governing blood flows, Eq 9, and of summary statistics such as the expectation (Eqs 4 and 10), the covariance (Eqs 5 and 11), the coefficient of variance, and the direction agreement rate (Eq 6). However, an analytical approach becomes unattainable when adopting a non-linear simulation model (for example, including the phase separation effect) or when utilizing other specific statistical distributions over boundary conditions. In such scenarios, uncertainty distributions governing blood flows could empirically be estimated by sampling multiple boundary condition configurations from their governing distribution and propagating these samples through the simulation model to estimate the resulting distribution governing blood flows. The following section outlines a more advanced data assimilation setup incorporating boundary condition uncertainty, observed or literature-derived data, and potentially also a non-linear simulation model incorporating the phase separation effect.

#### 2.4.3. Bayesian model calibration and uncertainty quantification in inverse modeling

Forward modeling is a relatively straightforward process of simulating or predicting a system’s state ***h***(***θ***) from known parameters ***θ***. Its counterpart, inverse modeling, is generally much more challenging[26, 45]. Inverse modeling involves assimilating model simulations with observed data to infer latent model states and the underlying but unknown parameters that most likely resulted in the observations. In the present context, inverse modeling would amount to inferring latent hemodynamic variables and the unknown pressure boundary conditions based on an incomplete and noisy set of measured or literature-derived blood flow rates or velocities. This process is inherently more complex due to the need to solve for multiple parameters that could potentially explain the observed data. Furthermore, the solution to an inverse problem is not always unique, and there may be multiple sets of parameters that could have produced the observations.

The Bayesian approach offers a statistically rigorous framework for confronting these challenges in inverse modeling[47]. It employs Bayes’ rule

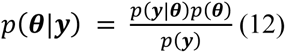

where *p*(***θ***|***y***) is the posterior probability of the parameters ***θ*** given the observations in ***y***, *p*(***y***|***θ***) is the likelihood which is the probability of the observations given the model simulations, *p*(***θ***) is the prior which is the probability of the parameters before seeing the observations, and *p*(***y***) is the marginal likelihood which is constant for a given set of observations. In the context of inverse modeling, the Bayesian approach allows for inferring the model’s latent states and parameters by comparing the model’s simulations with observed data. This comparison is quantified by the likelihood function, which measures how well the model’s simulations align with the observations. By utilizing probability distributions, the Bayesian approach not only enables the inference of the most probable values of the unknown latent variables and parameters ***θ***, but it also inherently quantifies the uncertainties associated with these estimates and naturally incorporates the range and likelihood of possible parameter values.

From a practical perspective, the Bayesian approach, as outlined above, requires the choice of likelihood and prior distributions, as indicated in Eq 12. Often, the posterior distribution in Eq 12 does not have an analytical solution which leads to the application of advanced techniques such as Markov Chain Monte Carlo methods[26, 47] for numerically computing the posterior distribution, along with summary statistics such as the expectation, Eq 4, the covariance, Eq 5, the coefficient of variance, and the direction agreement rate, Eq 6, in the present context. These practical considerations are further detailed in the forthcoming section on Experiment 2.

### 2.5. Microvascular networks

The analysis was conducted using extensive microvascular networks from the mouse somatosensory cortex[32, 50, 51]. Variants of these networks including blood flow simulations, subsequently made accessible by Schmid et al. [31, 52], and downloaded from https://doi.org/10.5281/zenodo.758632 and https://doi.org/10.5281/zenodo.5115639 under the CC BY 4.0 license, were used in the present study. The two networks, which are available in both data sources[31, 52] were considered. These networks, which are anatomically accurate, represent volumes of approximately 1.5-2.2 cubic mm from the pial surface to a depth of around 1.2 mm. Each network comprises thousands of vessel segments and nodes. Vessels were categorized into six classes based on their diameter and their connectivity with surface vessels of known classes[31]. Vessel classes included surface arterioles (SA), descending arterioles and arterioles (DA+A), capillaries (C), ascending venules and venules (AV+V), surface venules (SV), and unknown (UNK). A histogram-based upscaling approach was employed to slightly upscale the capillary diameters while preserving the hierarchical order of individual vessel diameters[31]. The two networks are visualized in Fig 1A.1 and 1B.1 and characteristic morphological parameters are summarized in Table 1 in S2 File.

**Fig 1.**
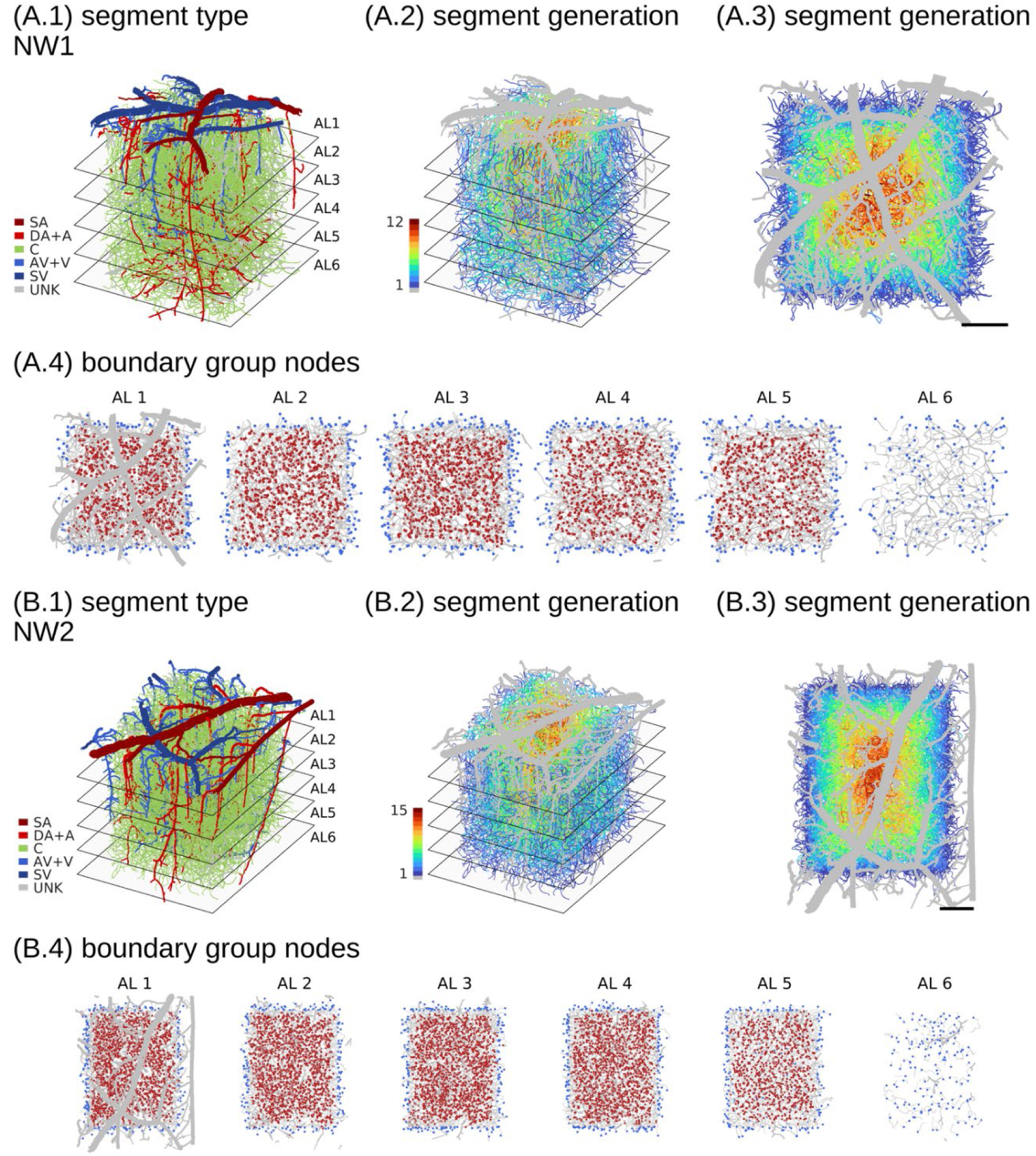
Segment types and segment generations for the two microvascular networks, NW1 and NW2. In (A.1, B.1), each segment is uniquely colored according to its type. In (A.2, B.2) and (A.3, B.3), the generation of capillary segments relative to network boundary nodes is color-coded, with non-capillaries shown in gray to emphasize capillary vessels. The first two columns present 3D (xyz) representations of the networks, while the third column provides 2D (xy) projections. Planes in the depth direction, visible in the first two columns, mark the boundaries between analysis layers (ALs), each separated by 200 µm. Scale bars in (A.3, B.3) represent 200 µm. In (A.4, B.4), boundary nodes of type C and UNK are colored blue, interior reference nodes are red, and vessel segments are gray. Some segments appear without both end-nodes since these nodes were located outside the respective analysis layers. SA: surface arteriole, DA: descending arteriole, A: arteriole, C: capillary, V: venule, AV: ascending venule, SV: surface venule, UNK: unknown.

**Table 1.**
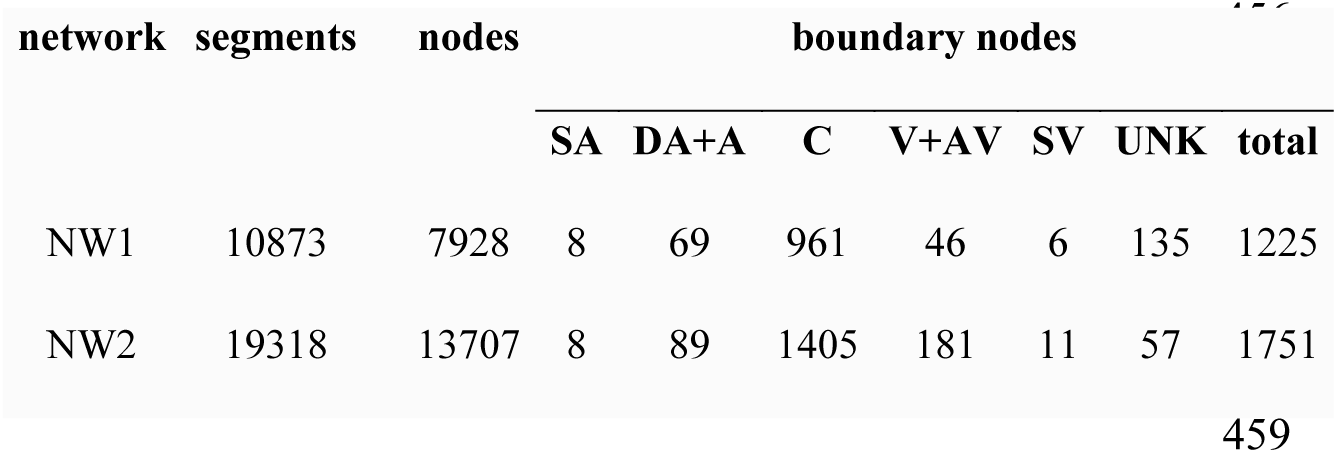
Segment and node information on the two microvascular networks (NW1 and NW2). Abbreviations: SA: surface arteriole, DA: descending arteriole, A: arteriole, C: capillary, V: venule, AV: ascending venule, SV: surface venule, UNK: unknown.

| network | segments | nodes | boundary nodes |  |  |  |  |  |  |
| --- | --- | --- | --- | --- | --- | --- | --- | --- | --- |
|  |  |  | SA | DA+A | C | V+AV | SV | UNK | total |
| NW1 | 10873 | 7928 | 8 | 69 | 961 | 46 | 6 | 135 | 1225 |
| NW2 | 19318 | 13707 | 8 | 89 | 1405 | 181 | 11 | 57 | 1751 |

Schmid et al. simulated the distribution of pressure, blood flow, RBC velocity, and hematocrit throughout the networks using a blood flow simulation model that tracks discrete RBCs[31]. This model incorporated an in-vitro formulation of blood’s effective viscosity and descriptions of the phase separation effect. Pressure boundary conditions were assigned using empirical diameter-dependent pressure values for surface arterioles and a uniform pressure for surface venules. For capillaries, pressure boundary conditions were assigned using a hierarchical approach in which the networks were embedded in a larger artificial microvascular network. The discrete nature of the simulation model leads to fluctuations in pressure, blood flow, and hematocrit over simulation run-time. Time-average of these parameters were considered for analysis and are available in the published data set. In the present study, these averages, denoted ref.Data, were used as a reference for evaluating the impact of boundary condition uncertainty on model simulations and in the quantitative assessment of the proposed adaptive method for setting pressure boundary conditions.

The networks, which essentially represent cubes cut from cortex, have numerous cut vessels on all six cube faces, except at the pial face. The latest version of the networks[52] features a somewhat simplified representation of vessel segments near network faces, and the original networks[31] were trimmed accordingly in the present study, with details provided in Fig 1 in S3 File. The number of boundary nodes for surface arterioles and surface venules is in the tens, but this number rapidly increases with depth, resulting in a large total number of boundary nodes: 1225 for the smaller network (NW1) and 1751 for the larger network (NW2), Table 1.

In the present study, a straightforward segment generation metric was further assigned to individual vessel segments. This metric, designed to be purely topological in nature, was defined in relation to boundary nodes and calculated using the following algorithm. Initially, a set of nodes was identified from all boundary nodes. Following this, a new set of nodes was identified which could be reached by propagating along segments connected to the initial set of nodes. These two node sets were then merged, and all segments with both their end-nodes contained in this combined set were assigned a generation level of one. A new iteration was then initialized from the combined set of nodes to identify segments with a generation level of two. This iterative procedure was repeated until a segment generation was assigned to all vessel segments, resulting in segment generations up to 12 (NW1) and 15 (NW2). The segment generations are visualized in Fig 1A.2/3 and 1B.2/3.

Groups of interior capillary nodes were identified to act as reference nodes in the proposed adaptive method for setting pressure boundary conditions. Each network was first divided into six analysis layers (ALs) along the depth direction, with each layer separated by 200 µm[31]. For AL 1 to 5, groups of reference nodes were then identified as the set of nodes that were end-nodes of capillary segments with a segment generation level of at least two to reduce the influence of boundary effects. The deepest analysis layer, AL 6, contained only a small number of candidate reference nodes. Therefore, no reference group was defined based on nodes in AL 6. Instead, the group of reference nodes in AL 5 was also utilized as reference nodes for boundary nodes in AL 6. Individual reference node groups contained 661 to 1071 (NW1) and 922 to 2287 (NW2) nodes, with details available in Table 2 in S2 File. The groups of reference nodes and associated boundary nodes are visualized in Fig 1A.4 and 1B.4.

Individual segments were assigned to an unique AL based their average endpoint depth, and hemodynamic metrics were analyzed for AL 1 to 5[31].

### 2.6. Experiments

#### 2.6.1. Experiment 1

The first numerical experiment was conducted to assess the impact of boundary condition uncertainty on model simulations and to quantitatively evaluate the proposed adaptive method for setting pressure boundary conditions. The experiment employed segment hematocrits and boundary node pressures from the reference dataset, ref.Data. To exclusively evaluate the impact of pressure boundary conditions, the following strategy was adopted. Hematocrit was not considered as a functional model variable in Experiment 1. Instead, hematocrit values from the reference data set were used to compute the effective viscosities of individual segments (after converting tube hematocrits to discharge hematocrits as described in Section 1.2.1 in S1 File). These viscosities were subsequently used to calculate the hydraulic resistances. These resistances, and consequently the matrix ***M*** in Eqn 1 and 2, were then kept constant throughout the simulations, enabling a direct assessment of the influence of boundary pressures parameterization across models. Consistent with the reference analysis[31], viscosities were calculated using an in-vitro viscosity formulation, plasma viscosity was set to 1.2, and no adjustment was made for differences in mean corpuscular volume (MCV) between human and mice.

Two models were examined and are referred to as *reference models* throughout the text. In the first model, ref.Vitro, model simulations were performed using node pressures from ref.Data as pressure boundary conditions for all boundary nodes. In the second model, ref.Vitro.ABC, node pressures from ref.Data were used as boundary pressures for vessels with types of SA, DA+A, AV+V, and SV, while boundary conditions for vessel types C and UNK were assigned using the proposed adaptive method. Specifically, the pressure boundary condition for a given boundary node (in a given AL) was assigned relative to reference nodes in the respective layer (Fig 1A.4 and 1B.4). Equal weighting coefficients were assigned to individual reference nodes, and the relative boundary pressures were thus defined relative to the average layer-wise reference node pressure. The relative boundary pressures *Δp*_*i*_ in Eq 3 were defined by sampling each of these i.i.d. from a Gaussian distribution with zero mean and a standard deviation of 1 mmHg. Thereby, heterogeneity in boundary pressures was induced by this random assignment of pressure deviances, while consistency between the layer-wise average boundary node pressure and interior reference node pressures was ensured at the same time.

The decision to use pressures from ref.Data for vessel types SA, DA+A, AV+V, and SV, and the adaptive method for vessel types C and UNK, mirrors a typical analysis scenario in which reference data for larger vessels SA, DA+A, AV+V, and SV would be available from the literature[30, 31, 33], albeit with some degree of uncertainty. In contrast, the pressures for the other vessels are subject to a higher level of uncertainty. Consequently, identical pressure boundary conditions was thus applied for vessel types SA, DA+A, AV+V, and SV in the ref.Vitro and ref.Vitro.ABC models, whereas boundary conditions for vessel types C and UNK differed across the two models.

The probabilistic approach to uncertainty analysis in forward models, Section 2.4.2, was adopted to examine the influence of pressure boundary condition uncertainty on blood flow uncertainty in the two models. Blood flow uncertainty was quantified by the direction agreement rate, Eq 6. A standard deviation of 2 mmHg was used to represent boundary pressure uncertainty. Uncertainty was exclusively applied to boundary vessels of types C and UNK to focus the analysis on how blood flow was influenced by uncertainties associated with boundary pressures that differed across the ref.Vitro and ref.Vitro.ABC models. In Experiment 2, a more advanced strategy was adopted which included boundary condition uncertainty for all vessel types.

#### 2.6.2. Experiment 2

In the second experiment, the adaptive method for pressure boundary conditions was integrated into our Bayesian calibration framework[26]. This integration aimed at inferring distributions over pressures at all boundary nodes and simultaneously at quantifying the impact of the resulting uncertainty on blood flow simulations.

The framework’s utility was assessed by implementing the blood flow simulation model with three different viscosity formulations, all of which are currently used in blood flow simulations in brain cortex[26, 29, 30, 34, 36]: an in-vitro viscosity formulation[40], cal.Vitro.ABC model, and two in-vivo viscosity formulations[41, 42], cal.Vivo.ABC and cal.Esl.ABC models. The three resulting model variants are referred to as *calibration models* in the text. Plasma viscosity was set at 1.2, and adjustments for the differences in MCV between humans, 92 fL,[1] and mice, 45 fL,[53] were included as described in Section 1.1 in S1 File. The phase-separation effect was additionally included to account for varying hematocrit throughout the networks. However, to reduce computational costs during the Bayesian calibration stage, a uniform discharge hematocrit of 40% was assigned to all segments[34, 49] and held constant during calibration. Bayesian calibration yielded samples from the posterior distributions governing the pressure boundary conditions, Eq 12, and subsets of these samples were subsequently propagated through non-linear simulation models incorporating the phase-separation effect using a uniform inlet discharge hematocrit of 40% at boundary nodes[34, 36]. The rationale behind the stepwise modeling approach is further discussed in the discussion section.

Measurements of blood flow rates or RBC velocities are not available for the two networks. Instead, the following approach was established to provide target velocities for arterioles and venules representing measurements obtainable from in-vivo experiments using current measurement techniques. Specifically, simulated RBC velocities, obtained from the reference data set, and in-vivo measurements in awake mice, obtained from literature[54], were used to create target RBC velocities for the Bayesian calibration. Analysis of vessel type and diameter as ordering parameters has revealed a statistically significant association between diameter and velocities in DAs and AVs[54]. Linear fits of this association were consequently obtained from RBC velocities, in vessel categories SA and DA+A (resp. SV and AV+V) collectively referred to as Art (resp. Ven), from AL 1 in the reference data set and in the literature data for vessels with diameters < 30 µm, matching the range of available in-vivo velocity measurements[54]. These fits were subsequently used to prescribe target velocities in Art and Ven with vessel diameters < 30 µm in AL 1. Additionally, target blood flow directions in SAs, DAs, AVs, and SVs were determined by iterating through these vessels starting from presumed inflow and outflow end-nodes of SAs and SVs, respectively. The linear velocity fits, along with the underlying data are shown in Fig 2 in S3 File, and segments assigned with target velocities and target flow directions are highlighted in Fig 3 S3 File.

**Fig. 2.**
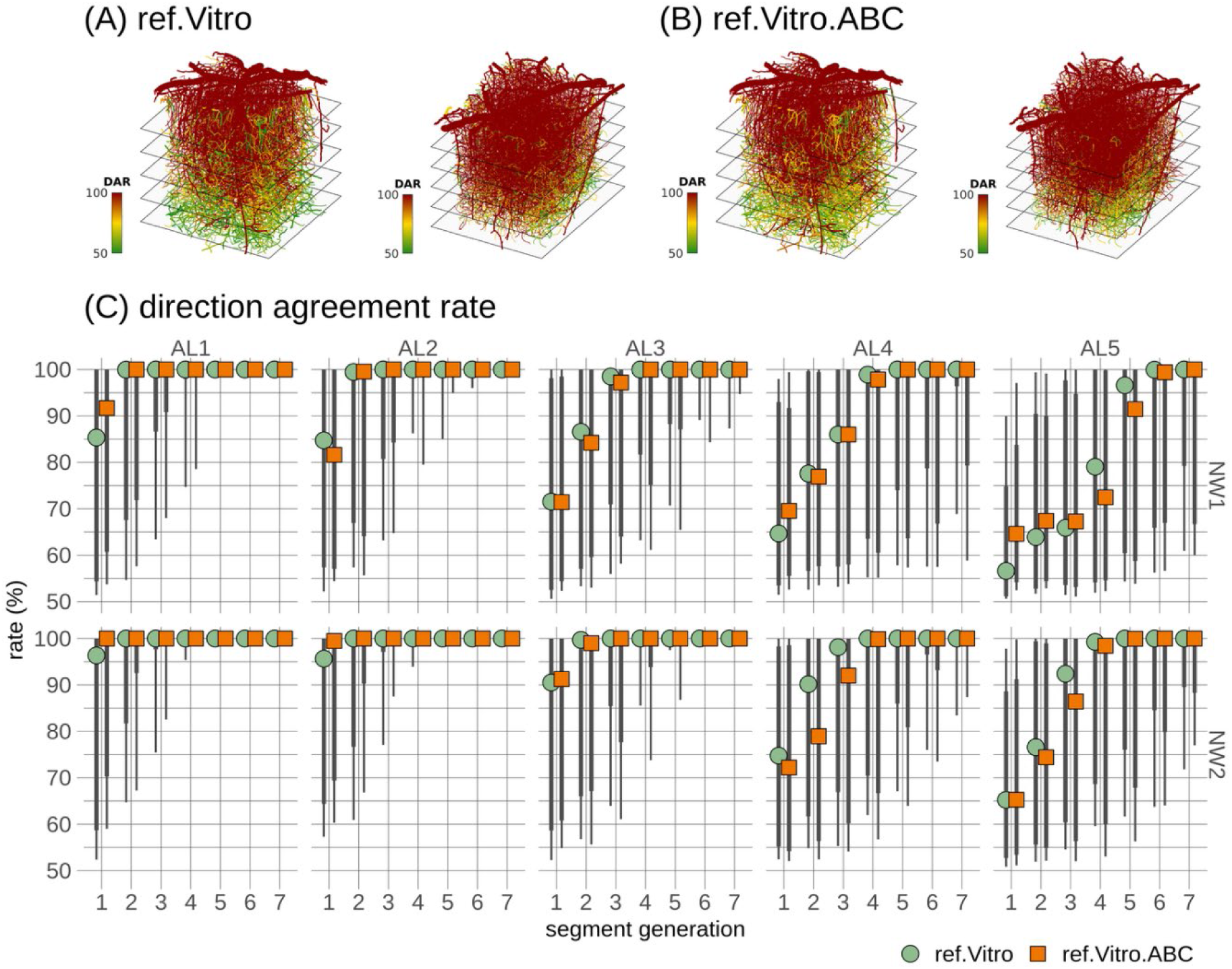
Quantitative analysis of flow direction uncertainty in the two reference models, ref.Vitro and ref.Vitro.ABC. In (A) and (B), each segment in the spatial representations of direction agreement rates (DAR) is uniquely colored according to its rate. A direction agreement rate of 100% corresponds to full agreement in blood flow direction, while a rate of 50% corresponds to complete disagreement. These rates are shown in the two models for both microvascular networks: one model with reference boundary pressures for all boundary segments (ref.Vitro), and the other with adaptive boundary pressures for capillary and unknown boundary segments (ref.Vitro.ABC). In (C), a summary of the direction agreement rates, grouped by network (NW1 and NW2), analysis layer (AL), and segment generation, is provided. Points denote medians across segments, while the thick and thin vertical lines represent [12.5 87.5] and [5 95] percentiles, respectively.

**Fig. 3.**
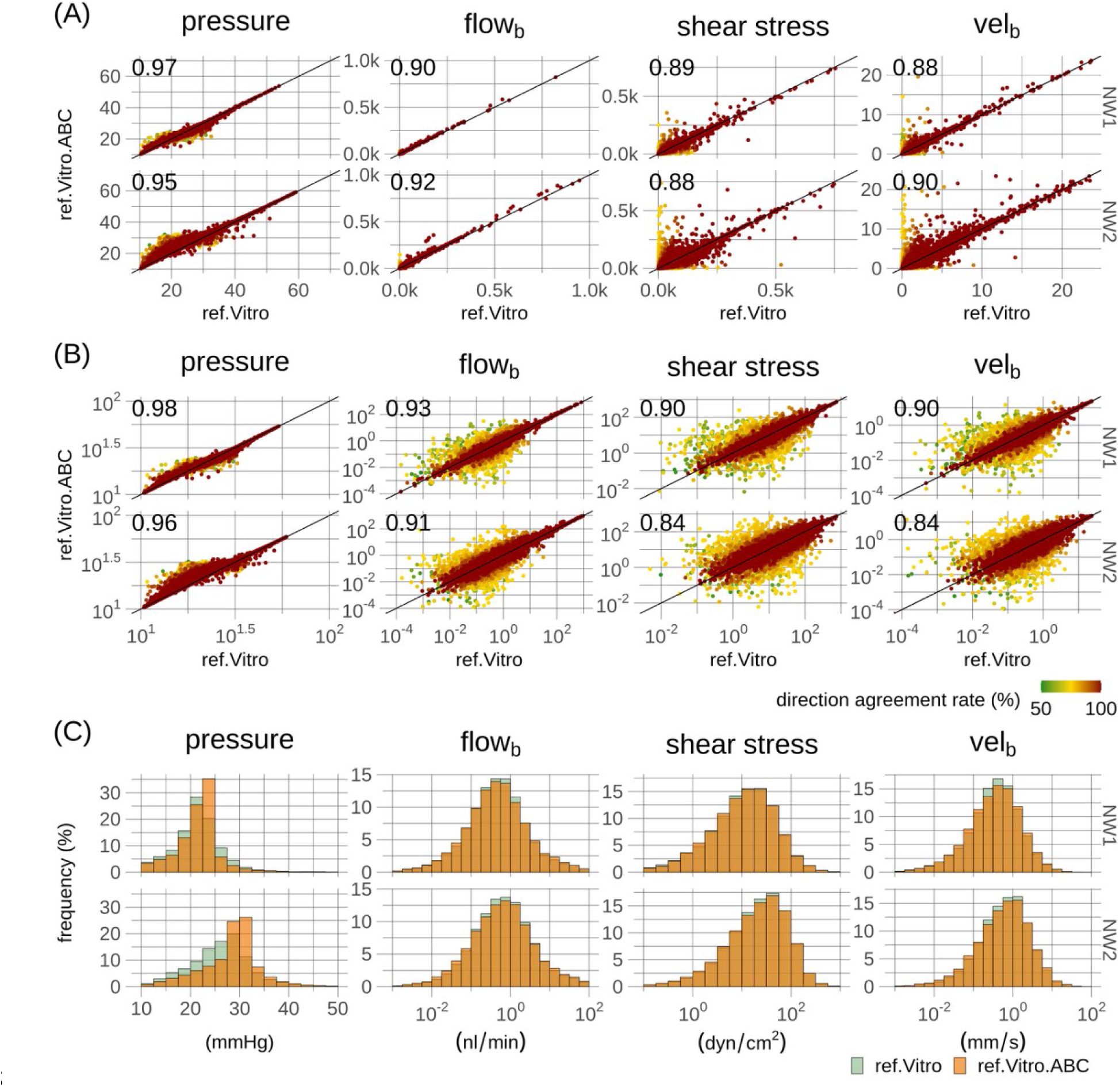
Quantitative comparison of hemodynamic metrics in the two reference models, ref.Vitro and ref.Vitro.ABC. In (A) and (B), each point in the scatter plots represents a vessel segment and is uniquely colored according to its direction agreement rate (DAR) averaged across the two reference models, ref.Vitro and ref.Vitro.ABC. For better visualization, scatter points are overlaid according to increasing DAR. The scatter plots are shown on a log-scale in (B). Black lines represent identity lines, and correlation coefficients, Spearman’s for (A) and Pearson for (B), are provided as inserts. Physical units of axes in (A) and (B) are consistent with the axis units of frequency histograms depicted in (C). Rows in (A-C) correspond to the two microvascular networks (NW1 and NW2).

Scaled and shifted beta distributions were used to describe uncertainties governing pressure boundary conditions[26, 39]. Broad symmetric distributions were applied to larger venules and arterioles (diameter threshold > 20 µm) with distribution support ranging from 5 to 15 mmHg for venules and 30 to 80 mmHg (resp. 50 to 100 mmHg) for arterioles in the cal.Vitro.ABC model (resp. cal.Vivo.ABC and cal.Esl.ABC models). Priors with higher pressure levels were adopted for the two models incorporating in-vivo viscosity formulations, due to the increased hydraulic resistance in these two models[41, 42]. Broad left-tailed distributions were applied to smaller arterioles with distribution support from 15 to 80 mmHg (resp. 15 to 100 mmHg) in the cal.Vitro.ABC model (resp. cal.Vivo.ABC and cal.Esl.ABC models). Symmetric distributions with support from 10 to 30 mmHg were applied to smaller venules. Relative pressure boundary conditions were applied at vessel types C and UNK with groups of interior reference nodes defined as in Experiment 1. The relative pressure boundary conditions, including their uncertainties, were modeled by symmetric distributions centered at 0 mmHg with distribution support from -8 to 8 mmHg and standard deviation of 2 mmHg, corresponding to the level of uncertainty used in the uncertainty quantification in Experiment 1. The prior distributions are shown in Fig 4 in S3 File, and mathematical details regarding likelihood functions and the prior distributions are further available in Sections 3.1 and 3.2 in S1 File.

**Fig. 4.**
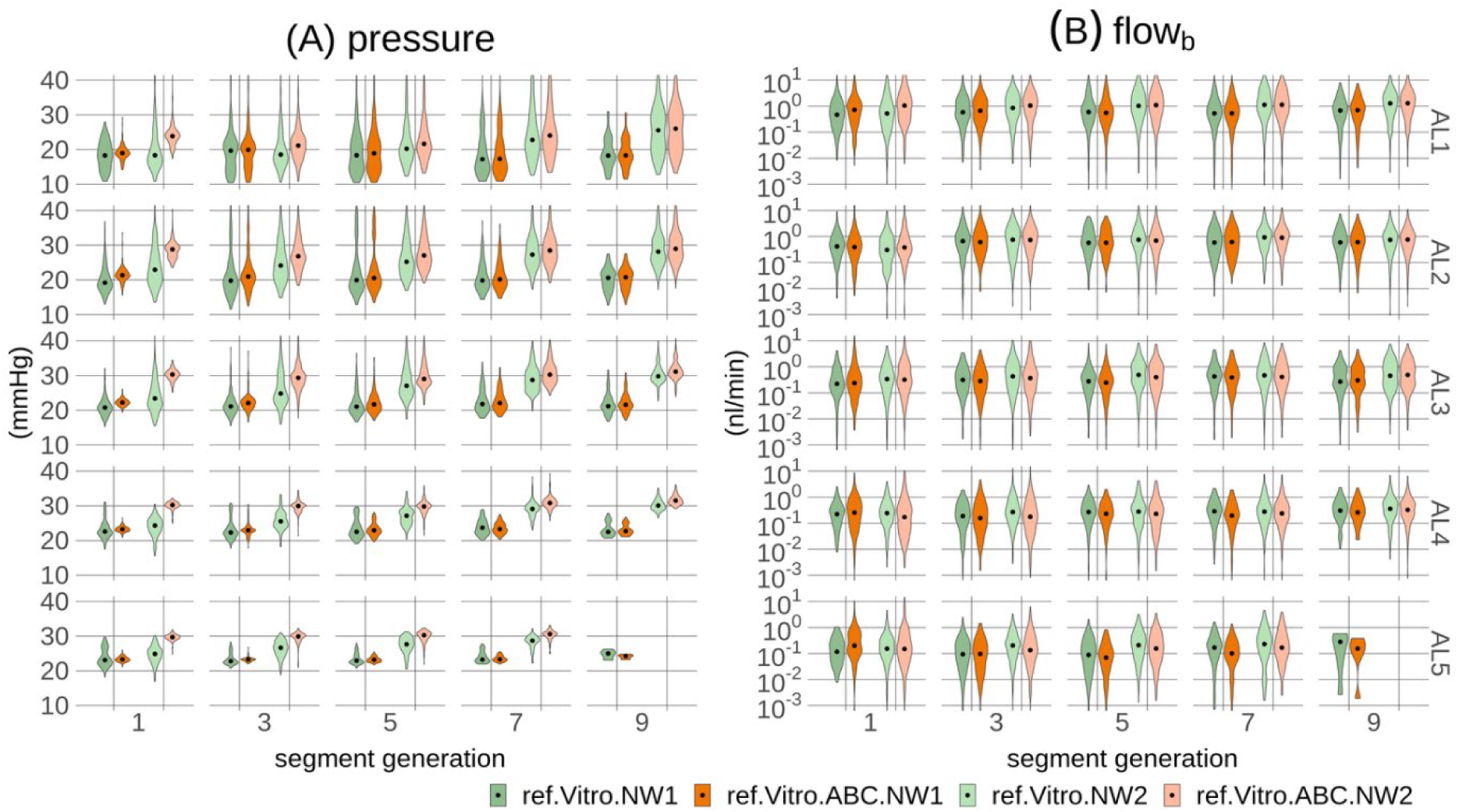
Depth-wise and generation-wise pressure profiles and blood flow profiles for capillary segments in the two reference models, ref.Vitro and ref.Vitro.ABC. In (A) and (B), different combinations of the two models (ref.Vitro and ref.Vitro.ABC) and the two microvascular networks (NW1 and NW2) are represented by unique colors. The plots are resolved by analysis layer (AL) in rows and segment generation in columns (see Fig 1 definitions of ALs and segment generation). Violins represent distributions over segments, and dots mark medians.

For Bayesian calibration, the DREAM(ZS) algorithm[26, 55–57] was used for parameter inference, with algorithmic settings detailed in Section 3.3 in S1 File. Five Markov Chain Monte Carlo (MCMC) chains were run in parallel, with each chain undergoing 2 million iterations, resulting in a total of 10 million MCMC iterations. The DREAM(ZS) algorithm produced an archive of 100,000 samples, representing a thinned history of the MCMC chain samples. The convergence of the sampled chains to a limiting distribution was assessed using the Gelman Rubin statistics using a threshold of 1.2 to declare convergence[26, 47, 57, 58]. A subset of 1,000 samples were retained for further analysis. The retained samples of pressure boundary conditions, one set of 1,000 samples for each of the three calibration models per network, were then used with the respective flow simulation models that now incorporated the phase-separation effect. This resulted in 1,000 simulations per model per network, each reflecting diverse boundary pressure configurations and variable segment hematocrit levels for the cal.Vitro.ABC, cal.Vivo.ABC, and cal.Esl.ABC models.

Hemodynamic metrics, for each segment, were summarized by averaging across the 1,000 model simulations. Similarly, uncertainty was quantified by the direction agreement rate calculated by the frequency at which the flow direction matched the predominant flow direction observed across the simulations.

## 3. Results

### 3.1. Experiment 1 – reference models

#### 3.1.1 Validation of reference simulations against reference data

Initially, the ref.Vitro model simulations were validated against the reference data, ref.Data, through a segment-wise comparison of the following key hemodynamic variables[34]: pressure, blood flow, shear stress, and blood flow velocity. Given that non-transformed variables typically span several orders of magnitude[3, 30, 31, 33, 34], potentially heavily influencing the Pearson correlation coefficient due to asymmetric distributions and potential outliers, Spearman’s rank correlation coefficient was used to quantify the monotonic relationship between non-transformed variables. For log-transformed variables[28], which mitigate the influence asymmetric distributions and potential outliers, the Pearson correlation coefficient was utilized. Strong correlations were observed for non-transformed variables, with a range from 0.98 to 1.00 (Spearman’s), as shown in Fig 5A in S3 File. Similar strong correlations were observed for log-transformed variables, ranging from 0.93 to 1.00 (Pearson), as shown in Fig 5B in S3 File. Additionally, there was a high congruence between frequency distributions, as shown in Fig 5C in S3 File, with the distribution variance between NW1 and NW2 in ref.Data being maintained in the ref.Vitro model simulations. Summary statistics governing the hemodynamic variables aggregated across the two networks and all vessel classes are available in Table 3 in S2 File, and more detailed summaries decomposed by vessel type and additionally by networks are available in Table 4 to 8 and 9 to 14 in S2 File, respectively. Overall, these results demonstrate that the ref.Vitro model simulations align closely with the hemodynamic metrics from ref.Data and thus provide a valid reference for subsequent analyses in the present study.

**Fig. 5.**
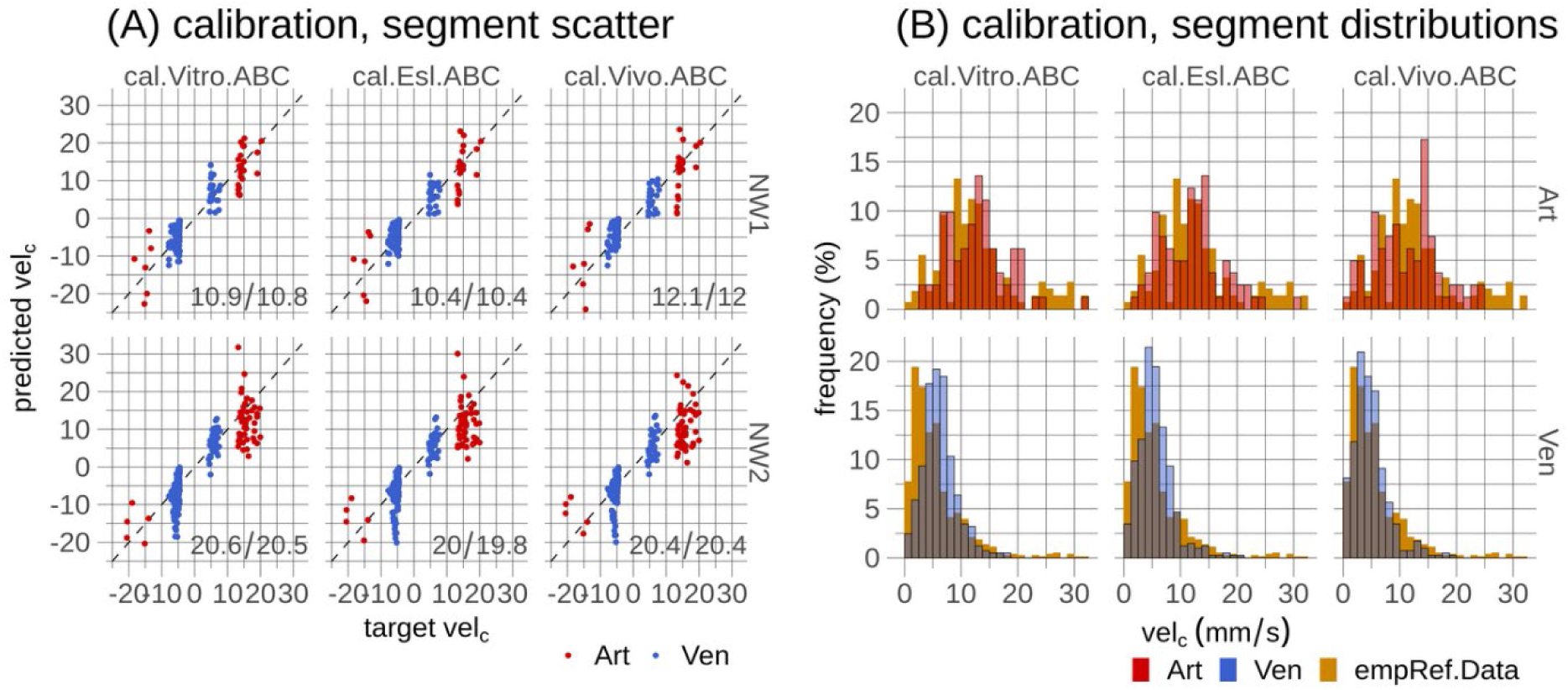
Bayesian model calibration in the three calibration models, cal.Vitro.ABC, cal.Esl.ABC, and cal.Vivo.ABC. In (A), scatter plots comparing predicted red blood cell velocities with target velocities are shown. Points represent individual segments, dashed lines represent identity lines, and mean squared errors are shown as inserts. The three calibration models are shown in columns, and the two microvascular networks (NW1 and NW2) in rows. Segments where target velocities were used in model calibration are shown in Fig 3 in S3 File. In (B), frequency histograms of predicted velocities in (A) are shown with vessel types in rows. These histograms are compared to the literature and reference data, empRef.Data, used to create diameter dependent target velocities. See Section 2.6.2 and Fig 2 and 3 in S3File for further information about this data and the creation of target velocities.

#### 3.1.2. Uncertainty quantification

After validating the ref.Vitro model simulations against ref.Data, the impact of pressure boundary condition uncertainty on blood flow uncertainty was evaluated in the ref.Vitro and ref.Vitro.ABC models. The hydraulic conductance of individual segments was identical across the two models, and pressure boundary conditions for boundary segments classified as SA, DA+A, AV+V, and SV were also identical across. However, the two models differed in the application of the adaptive method for boundary pressures in the ref.Vitro.ABC model at segments categorized as C and UNK. The spatial distributions of blood flow uncertainty, quantified by the direction agreement rate metric, are shown in Fig 2A and 2B. The direction agreement rates ranged from 50% to 100%, displaying considerable spatial heterogeneity. This spatial heterogeneity was further quantitatively analyzed by grouping segments based on their respective analysis layer and segment generation, as shown in Fig 2C. An increase in direction agreement rate with increasing segment generation was observed across all analysis layers. Additionally, a depth-dependent influence was observed, with only the first few segment generations being markedly affected by blood flow direction uncertainty in AL1, while uncertainty escalated with depth in higher generation levels. The ref.Vitro.ABC model exhibited marginally lower median values with increasing depth, but these minor differences should be considered in light of the extensive distributions represented by the vertical lines in Fig 2C and further detailed in Fig 6A in S3 File. Consequently, a consistent pattern of spatial heterogeneity in blood flow uncertainty was observed between the two reference models across the two microvascular networks.

**Fig 6.**
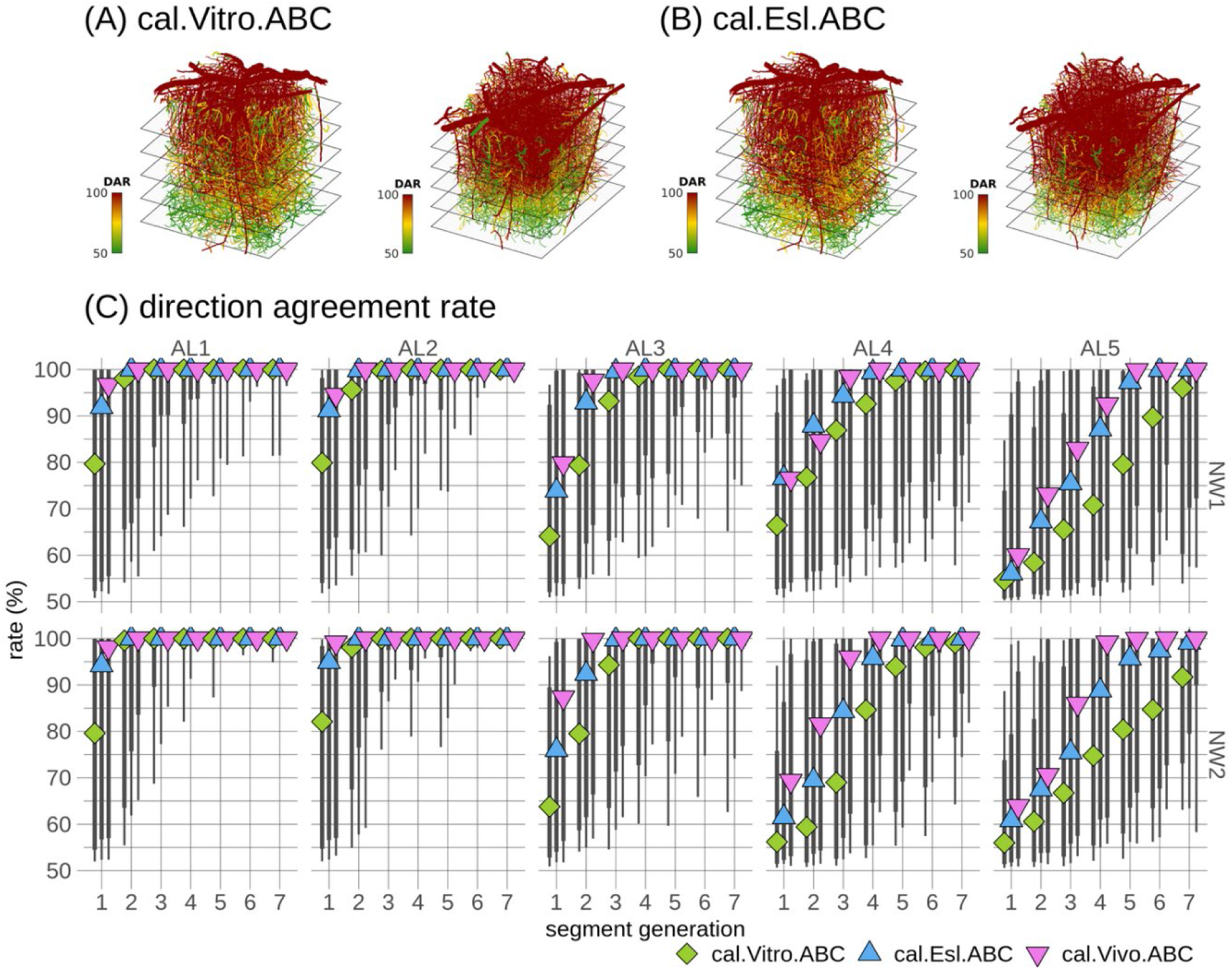
Quantitative analysis of flow direction uncertainty in the three calibration models, cal.Vitro.ABC, cal.Esl.ABC, and cal.Vivo.ABC. Same format as in Fig 2 but showing the cal.Vitro.ABC and cal.Esl.ABC models in (A) and (B) and all three calibration models in (C). The spatial representation of direction agreement rate (DAR) for the cal.Vivo.ABC model is similar to (B) and is available in Fig 13 in S3 File.

#### 3.1.3. Hemodynamic predictions

Segment-wise comparisons of the ref.Vitro and ref.Vitro.ABC models’ predictions of segment pressure, blood flow, shear stress, and blood flow velocity are shown in Fig 3A and 3B. Strong correlations were observed for the non-transformed variables, with a range of 0.88 to 0.97 (Spearman’s), as well as for the log-transformed variables with a range from 0.84 to 0.98 (Pearson). Segments governed by a low direction agreement rate exhibited the strongest scatter between the two models. A detailed analysis of the relationship between direction agreement rate (within model) and flow direction consensus (between models), over various direction agreement rate thresholds, is provided in Fig 7 in S3 File. It was observed that although the number of super-threshold segments naturally decreased with increased threshold, a high number of super-threshold segments remained, while the number of segments lacking flow direction consensus between the two models diminished. Taken together, these findings demonstrates robust consistency in hemodynamic predictions between the ref.Vitro and ref.Vitro.ABC models, and further suggest that uncertainty analysis offers an objective means for identifying a set of segments that exhibit a strong correlation between the hemodynamic variables in the two models.

**Fig 7.**
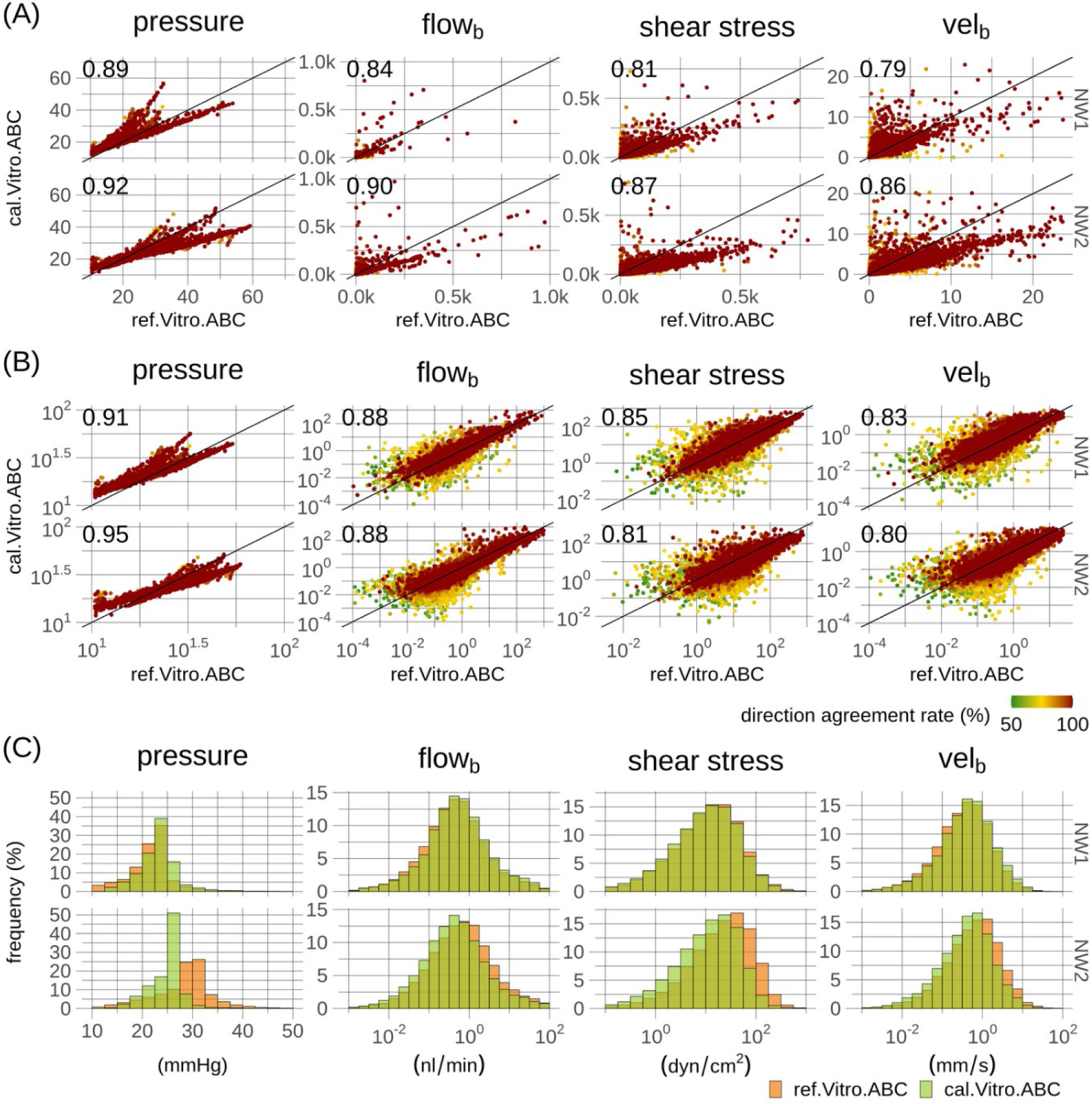
Quantitative comparison of hemodynamic metrics of the ref.Vitro.ABC and cal.Vitro.ABC model. Same format as Fig 3 but with a comparison of the ref.Vitro.ABC and cal.Vitro.ABC models.

Fig 3C shows frequency distributions of the hemodynamic variables, which demonstrate a notable congruence between the two models. The ref.Vitro.ABC model effectively captured the variations in blood flow rates, shear stress, and velocity across the two networks as observed for the ref.Vitro model. The ref.Vitro.ABC model exhibited slightly more compact pressure distributions, in particular for NW2, than the ref.Vitro model. A detailed examination of this phenomenon across ALs and segment generations, shown in Fig 4A, revealed consistent distribution patterns within segment subgroups, with differences in mean pressures mainly occurring in the lower segment generations. Specifically, NW2 exhibited elevated pressures compared to NW1 in higher generations in both models. The ref.Vitro model, however, showed a decline in pressure at lower generations, aligning with NW1 levels, whereas the ref.Vitro.ABC model maintained a relatively uniform pressure across generations. In interpreting these differences it should be noted that the boundary conditions for the ref.Vitro model were derived from simulations incorporating the networks within a larger synthetic network, leading to anticipated similar pressure levels for the two networks near their boundaries. The ref.Vitro.ABC model incorporated the adaptive approach, setting pressure boundary conditions relative to interior reference node pressure, anticipating more uniform pressure profiles. Despite these nuanced pressure variations, comparable pressure gradients were noted (Fig 6B in S3 File), and blood flow simulations indeed showed a strong agreement, as shown in Fig 3A to 3C and Fig 4B, and in the summary metrics in Table 3 to 14 in S2 File. Furthermore, the pressure variations between the two models were confined to the combinations of ALs and segment generations governed by the most uncertainty, Fig 2C and 4A.

### 3.2. Experiment 2 – Bayesian calibration

Following Experiment 1, which established a strong correspondence between the ref.Vitro and ref.Vitro.ABC models and thereby supported the validity of the proposed adaptive method for pressure boundary conditions, the method was incorporated into our Bayesian calibration framework. This approach treated all boundary pressures as variables with inherent uncertainty and incorporated the adaptive method for pressure boundary conditions to boundary vessels with categories C and UNK as in Experiment 1. Target flow directions and RBC velocities, Fig 2 in S3 File, were used in specific arterioles and venules, Fig 3 in S3 File, and three distinct viscosity formulations were adopted. Convergence of the MCMC sampled chains to a limiting distribution was assessed by the Gelman Rubin *R*_*c*_ statistics, which confirmed convergence for all three models. Sets of samples of inferred pressure boundary conditions were subsequently utilized in models that incorporated the phase-separation effect, to account for non-uniform hematocrit distribution.

#### 3.2.1. Hematocrit iterations – model convergence

Examples of blood flow changes and hematocrit changes across hematocrit iterations, for the three calibration models, are shown in Fig 8 to 10 in S3 File and discussed in Section 2 in S1 File. Frequency histograms showing the required number of iterations needed for convergence, for the 1,000 distinct pressure boundary configurations per calibration model in each network, are presented in Fig 11 in S3 File and summarized in Table 15 in S2 File. The cal.Vitro.ABC model exhibited the quickest convergence, necessitating 18/17, NW1/NW2, iterations (median), followed by the cal.Esl.ABC model (31/33 iterations), while the cal.Vivo.ABC model demanded the greatest number of iterations (56/62 iterations), as shown in Fig 11 in S3 File and Table 15 in S2 File. Permitting a minority of segments (ten) to surpass the convergence criterion substantially decreased the iteration count to 15/15, 23/24, and 31/32, for cal.Vitro.ABC, cal.Esl.ABC, and cal.Vivo.ABC, respectively, with negligible impact on the resultant blood flow and hematocrit estimates (Fig 12 in S3 File). Taken together, the iterative procedure exhibited good convergence efficacy across all three model variants in both networks when exposed to the pressure boundary conditions inferred by our Bayesian calibration framework.

**Fig 8.**
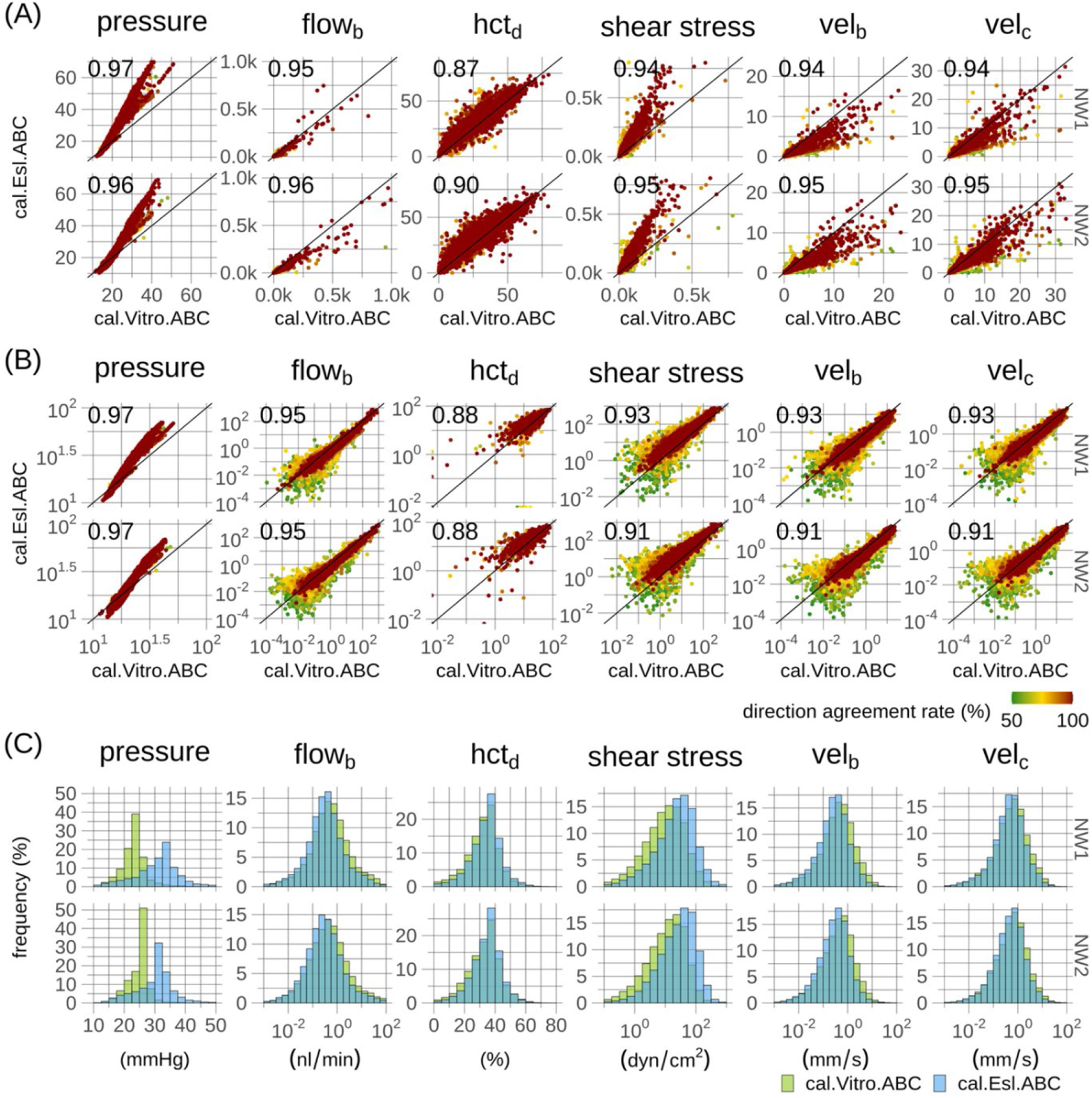
Quantitative comparison of hemodynamic metrics in two calibration models, cal.Vitro.ABC and cal.Esl.ABC. Same format as Fig 3 but with a comparison of the cal.Vitro.ABC and cal.ESL.ABC models. Hematocrits and red blood cell velocities are additionally compared, since these were considered as functional variables in both models.

#### 3.2.1. Calibration fits

Segment-wise comparisons of target velocities against predicted velocities are presented in Fig 5A. The three viscosity formulations yielded similar performance, as quantified by the mean squared velocity error, with similar errors observed in vessel categories Art and Ven. Scatter points tend to form clusters along the horizontal axis (target velocity) since vessels governed by target velocities have similar diameters, Fig 3 in S3 File. Points within these clusters exhibited scatter along the vertical axis (predicted velocity), resulting from the diameter-dependent target velocities being adopted and agrees with the scatter observed for a given diameter in the data underlying the target values, as seen in Fig 2 in S3 File. Frequency distributions governing the predicted velocities and the data underlying the target velocities are shown in Fig 5B. High congruence between the distributions is observed for all three models, and the models captured well the distributional differences between Art and Ven observed in the data underlying target velocities.

#### 3.2.2. Uncertainty quantification

The spatial distributions of blood flow uncertainty, as quantified by the direction agreement metric, for the three calibration models are shown in Fig 6A and 6B and Fig 13 in S3 File. Consistent with observations in the reference models, Fig 2A and 2B, substantial heterogeneity in segment-wise uncertainty was observed, with the direction agreement rate characterized by both depth-dependent and generation-dependent influences. Considerable heterogeneity was also observed among vessels within a given depth and generation group as shown in Fig 6C. The medians governing the depth and generation sub-groups of the cal.Vivo.ABC and cal.Esl.ABC models followed each other and exhibited escalating uncertainty with depth, in agreement with the observation in the reference models, Fig 2C. The cal.Vitro.ABC model exhibited a similar pattern but with greater uncertainties for deeper analysis layers, AL3 to AL5, compared to the two other calibration models, Fig 6C, and to the ref.Vitro and ref.Vitro.ABC models, Fig 2C. When interpreting these differences, the extensive distributions, indicated by vertical lines in Fig 6C and further detailed in Fig 14A in S3 File, should be considered. Additionally, identical target velocities and identical distributions, representing boundary conditions uncertainties for vessel types C and UNK, were used in all three calibration models. However, the ref.Vitro.ABC model incorporated the in-vivo viscosity formulation, resulting in reduced hydraulic resistance compared to the two other calibration models incorporating in-vivo viscosity formulations. Given that the same target velocities were used, and that reduced pressures in the cal.Vitro.ABC were expected and observed, Table 3 to 14 in S2 File, the stronger influence of boundary condition uncertainty on blood flow uncertainty in the cal.Vitro.ABC model was consequently expected.

#### 3.2.3. Hemodynamic predictions

To establish a link between Experiments 1 and 2, the ref.Vitro.ABC and cal.Vitro.ABC models’ predictions of pressure, blood flow rate, shear stress and blood flow velocity were compared at a segment-wise level as shown in Fig 7A and 7B. Strong correlations were observed for the non-transformed variables, ranging from 0.79 to 0.92 (Spearman’s), as well as for log-transformed variables with a range from 0.81 to 0.95 (Pearson). Categorizing segments according to their blood flow direction agreement rates, as in Experiment 1 Fig 3, demonstrated most scatter among segments with low direction agreement rate and least scatter among segments high direction agreement rate in agreement with the result in the reference models, Fig 3A and 3B. Frequency distributions, shown in Fig 7C, demonstrated high congruence between the ref.Vitro.ABC and cal.Vitro.ABC models in NW1, whereas the cal.Vitro.ABC model exhibited slightly left-shifted distributions in NW2. On the other hand, high congruence was observed between the two networks in the cal.Vivo.ABC whereas larger differences between the two networks were observed for the ref.Vivo.ABC model. The reason for the higher congruence between the two networks for the calibration model is that the same empirical description of target RBC velocities were utilized for both networks.

Segment-wise comparisons between two of the calibration models, cal.Vitro.ABC and cal.Esl.ABC, and corresponding frequency histograms are shown in Fig 8A to 8C. This comparison also included hematocrits and RBC velocities since these variables were considered as functional variables in both models. Although a uniform hematocrit of 40% was imposed at inlets, hematocrit quickly became heterogeneous as inflowing blood encountered bifurcations, resulting in large hematocrit dispersion as seen in Fig 8. Strong correlations were observed for both non-transformed variables from 0.87 to 0.97 (Spearman’s) and log-transformed variables from 0.88 to 0.97 (Pearson). A high congruence was observed for the frequency histograms (Fig 8C) across the two networks and across the two models, except for the pressure and shear stress that exhibited left shifted distributions for the cal.Vitro.ABC model. However, despite these shifted distributions, which were expected due to the reduced hydraulic resistance in the cal.Vitro.ABC model, a strong segment-wise correlation of 0.96 to 0.97 (resp. 0.91 to 0.95) was observed for pressure (resp. shear stress), Fig 8A and 8B, demonstrating a strong consistency in the hierarchy governing segment pressures across these two models. A comparison between the cal.Vitro.ABC and cal.Vivo.ABC models is shown in Fig 15 in S3 File and demonstrated a strong between-model consistency, like the comparison in Fig 8 but with a slightly increased scatter. A comparison between the two in-vivo models, cal.Esl.ABC and cal.Vivo.ABC, is shown in Fig 16 in S3 File demonstrated, on the other hand, the strongest histogram congruence between any pair of calibration models being compared. Summary statistics governing all hemodynamic metrics in all models are available in Table 3 to 14 in S2 File.

### 3.3. Path-based analysis of pressure profiles and capillary flow patterns across different viscosity models and pressure boundary conditions

So far, the quantitative comparisons established a good correspondence between the two reference models, between the three calibration models, and between the reference and calibration models. Depth-dependent pressure profiles and layer-wise capillary RBC flow patterns have previously been demonstrated[31]. A corresponding path-based analysis was conducted to evaluate the extent to which the proposed adaptive method for pressure boundary conditions reproduces these phenomena, and to examine whether the phenomena generalize across the three viscosity models including in-vivo viscosity formulations. Specifically, flow paths between SA and SV boundary nodes were tracked for all five models. The two reference models each provided a single simulation in each network, and a single set of paths was therefore obtained in each of these models. The calibration models each provided 1,000 simulations in each network. In the calibration models, path-tracking was therefore first performed on individual simulations, and the resulting sets of paths were then aggregated across simulations. Paths along which blood followed the vessel sequence SA→DA+A→C→AV+V→SV were retained, with up to two subsequent deviating transitions permitted and vessels with type UNK neglected, to render the path selection criteria more robust to vessel labeling errors[31].

Layer-wise pressure profiles are shown in Fig 9A.1. Depth-dependent pressure profiles were observed in all models in agreement with previously reported results[31]. A strong quantitative consistency was observed between the two reference models and between these models and previously reported results. Close to the pial surface (AL 1), the largest pressure drop occurs across vessels categorized as capillaries. Their contribution to the total pressure drop decreases with increasing cortical depth. On the other hand, the pressure drop across arterioles increases with depth and exceeds the capillary pressure drop around AL2 to AL3. The pressure drop in venules also increase with depth, whereas the contribution from surface vessels was minor. Pressure profiles observed for the cal.Vitro.ABC model are in quantitative agreement with the reference models but with lower magnitude, in agreement with the more compact pressure distribution in this model, Fig 7 and Table 3 to 14 in S2 File). The in-vivo models, cal.Esl.ABC and cal.Vivo.ABC, also provided depth-dependent profiles in qualitative agreement with the reference models, but with larger pressure drops than cal.Vitro.ABC consistent with their higher pressure levels and broader pressure distributions, Fig 8 and Table 3 to 14 in S2 File. Uncertainty analysis, Fig 9A.2, revealed that a high level of blood flow stability, as quantified by direction agreement rate, was maintained along the flow paths underlying the pressure profiles, Fig 9A.1, with slight decreases in direction agreement rate with increasing dept in agreement with Fig 2 and 6. Layer-wise capillary flow patterns were examined by considering the cortical depth of capillary start points and capillary end points along individual flow paths. Fig 9B.1 shows a strong correlation between the cortical depths of capillary endpoints in agreement with previous findings[31], and demonstrate that this phenomenon generalize to in-vivo viscosity formulations. Furthermore, the uncertainty analysis, Fig 9B.2, demonstrates a high average direction agreement rates for segments along individual paths, supporting the reliability of the depth-wise correlations.

**Fig 9.**
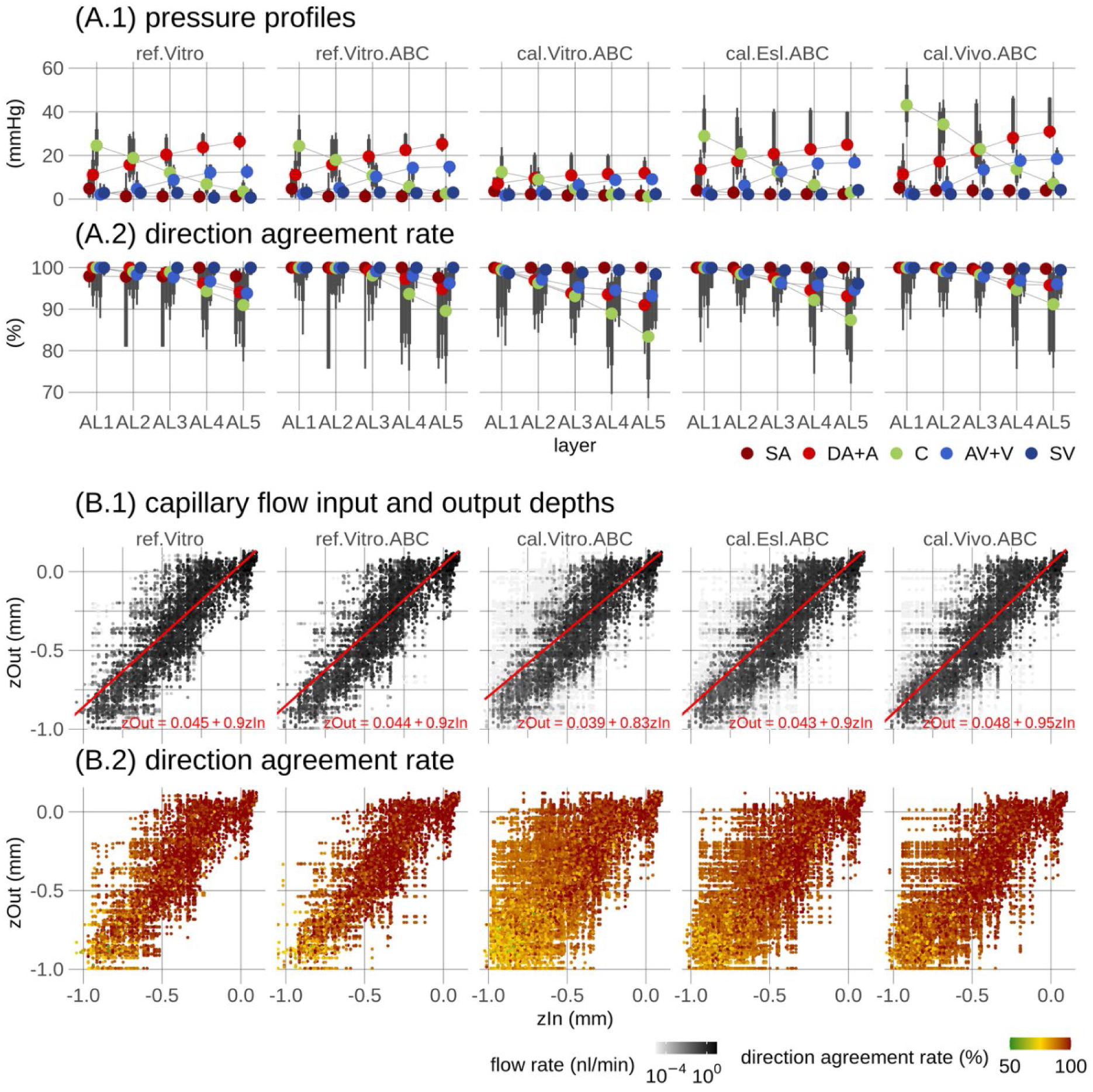
Pressure profiles and capillary input and output depths along pathways of blood flow. (A.1) shows pressure drop profiles for each vessel type are shown. The paths of blood flow, which were tracked from the end nodes of surface arterioles to surface venules, are grouped by analysis layers (ALs) based on the depth of the first capillary segment along the paths. The columns display the two reference models and the three calibration models. Points denote medians across paths, while the thick and thin vertical lines represent [12.5 87.5] and [5 95] percentiles, respectively. (A.2) shows a corresponding uncertainty analysis, with uncertainty quantified by average direction agreement rate of segments across respective parts of individual flow paths. In (B.1), the depths of capillary inputs (zIn) and outputs (zOut) are presented. Individual paths are represented by points, with the intensity of the greyscale color indicating the blood flow rate. For better visualization, colors are displayed on a log-scale, and points are ordered according to intensity. Inserts provide linear fits and their corresponding equations. The summary statistics in (A.1 and A.2) and the fits in (B.1) were computed by weighting individual paths according to their blood flow. (B.2) shows direction agreement rates corresponding to points in (B.1) using the same point ordering. SA: surface arteriole, DA: descending arteriole, A: arteriole, C: capillary, V: venule, AV: ascending venule, SV: surface venule.

Results provided in Fig 9A and 9B represent metrics that were weighted according to path blood flow rate, to facilitate a direct comparison across all five models, as hematocrit was only considered as a functional variable in the three calibration models. Fig 17 in S3 File provides complementary results based on weighting according to RBC flow rate in the three calibration models, and strong consistency between the two weighting strategies was observed.

## 4. Discussion

### 4.1. Network boundary conditions

Blood flow simulations in microvascular networks are heavily influenced by the selection of boundary conditions making this task a persistent challenge that requires considerable attention[26, 28–31, 33–38]. This work proposes an adaptive method for pressure boundary conditions, which, to the best of the author’s knowledge, has not been previously considered. The rationale behind the method is its reproducing property, ensuring that statistical properties of boundary nodes mirror those of interior reference nodes while maintaining flexibility at individual boundary nodes in terms of relative pressure deviations, Section 2.3.1. The method inherently adapts its reference pressure levels and thereby eliminates the need for declaration or iterative estimation of these levels when exposed to various microvascular networks with differences in vascular topology and vessel morphology, for example. The method is simple to implement, as it can be directly incorporated into the system of equations governing the blood flow model, Section 2.3.2. The method requires no additional iterations of consecutively solving the system of governing equations, as the reference pressure levels inherently adapt through the method’s incorporation into these equations.

The proposed adaptive method for pressure boundary conditions follows a similar spirit as previous methods prescribing an adjusted but constant pressure at capillary boundary nodes[30] and recently by prescribing pressures in cut arterioles and venules as equal to the mean pressure of the respective vessel classes of equal diameter at the same depth in the network[36]. Recent techniques have further focused on modeling heterogeneous, depth-dependent boundary conditions. This is achieved by implanting microvascular networks into a larger artificial network designed to mimic the structural properties of the actual network[31], or by eliminating capillary boundary nodes by connecting segments located at opposite network faces[36]. Instead of eliminating boundary nodes and their corresponding boundary conditions, the proposed method maintains these unknown variables but re-references them relative to sets of interior nodes. This ensures self-consistency in node pressures while not allowing blood trajectories to exit one domain boundary and re-enter at the opposite domain boundary, reflecting the unknown history of blood entering the domain[31]. While the proposed method eliminates the need for designing large synthetic embedding networks, it introduces the additional requirements of defining interior reference nodes, defining weighting coefficients, and defining relative pressure deviations. A reasonable strategy is to define reference nodes based on some similarity measures, such as node type, morphology, or topology. Similarly, defining weighting coefficients to yield the average reference node pressure, as done in this study, is a reasonable and pragmatic first strategy. Maintaining boundary conditions as free parameters in terms of relative pressure deviations is a desirable and unique feature of the proposed adaptive method as it acknowledges the inherent uncertainty of these boundary conditions, and it facilitates the method’s incorporation into data assimilation contexts as done in Experiment 2.

The adaptive method for pressure boundary conditions was exposed to two modeling scenarios. Experiment 1 represents a typical in-silico or forward modeling scenario. In the ref.Vitro.ABC model, pressure values were imposed at arterioles and venules (418 nodes in total, Table 1), whereas the adaptive method for pressure boundary conditions was applied to vessel types of C and UNK (2558 nodes in total, Table 1). Heterogeneity across the relative boundary conditions was incorporated by sampling these from random distributions. Experiment 2, on the other hand, represents a typical data assimilation scenario where the three calibration models (cal.Vitro.ABC, cal.Esl.ABC, and cal.Vivo.ABC) were combined with literature-derived RBC velocities to facilitate inference over the unknown boundary conditions and latent hemodynamic variables. To establish a link between Experiments 1 and 2, the adaptive method was also applied to vessel types of C and UNK in Experiment 2, whereas the remaining boundary pressures were considered as conventional uncertain boundary pressures in the three calibration models. It would, however, be straightforward to apply the adaptive method also to cut arterioles and venules in Experiment 2. In this case, additional depth-dependent groups of reference nodes could be defined for the respective segment types, and relative pressure boundary conditions could then be defined relative to these groups. An important feature of the adaptive method in this context, which distinguishes it from previous approaches[30, 36], is the application of the relative pressure deviations. For example, if information about flow directions in deeper cut arterioles and venules is incorporated into a data assimilation analysis, the model flexibility induced by the relative pressure deviations will allow the inferred boundary conditions to adapt to maintain agreement between the simulated and observed flow directions[26].

Guided by previous research, demonstrating decreased pressure with cortical depth[29, 31], the interior reference nodes were partitioned into sets according to their cortical depth and individual boundary nodes were matched with these sets according to their respective depth. Another approach could have been to use identical sets of reference nodes for all boundary nodes governed by the adaptive method, and instead incorporate systematic depth dependence through the relative pressure deviations. However, such an approach hinges on specific knowledge of the functional form governing pressure variations across depth. Alternatively, identical sets of interior reference nodes could be used for all relative pressure boundary conditions, whereas the individual weighting coefficients *w* in Eq 3 instead could vary according to the depth difference between a respective boundary node and its reference nodes. However, this approach could lead to reduced sparsity leading to a substantial increase in the computation effort needed to solve the system of equations in Eq 2. On the other hand, the approach taken in the present study preserved sparsity, as described in Section 2.3.3, and thereby maintained the required computational effort at a low level. As a result, this sparsity-preserving strategy guarantees that the adaptive method remains scalable when applied to large microvascular networks with a massive number of boundary conditions and interior reference nodes. The validity of the approach of utilizing groups of interior reference nodes is supported by the strong consistencies with simulations based on boundary pressures from the reference data set in hemodynamic variables (Fig 3 and Table 3 to 14 in S2 File), in the uncertainty profiles (Fig 2), and in the depth-dependent pressure profiles and layer-wise capillary blood flow profiles (Fig 9).

### 4.2. Flow simulations

To connect the findings of this study with previous research, a methodical approach was adopted, gradually increasing the complexity of the models. Initially, two reference models, ref.Vitro and ref.Vitro.ABC, were considered. Hydraulic conductance was calculated from hematocrit in reference data and kept consistent across both models, allowing for an exclusive evaluation of the proposed adaptive method for pressure boundary conditions against simulations based on pressure boundary conditions from reference data, Fig 3 and Table 3 to 14 in S2 File. The analysis demonstrated strong consistency between the two models, providing evidence that the adaptive method is suitable for defining pressure boundary conditions in extensive microvascular networks.

Once the usefulness of the adaptive approach was established, it was incorporated into our Bayesian calibration framework for a more complex analysis. Distributions over pressures at all boundary nodes were inferred by calibrating the simulation models against target RBC velocities and presumed flow directions in a subset of arterioles and venules. A two-step approach was adopted to reduce computational demand. During the Bayesian calibration stage, a uniform hematocrit of 40% was used for all vessel segments, aligning with previous studies that also adopted a uniform hematocrit across vessel segments[27, 34, 49]. The simplification utilizing uniform hematocrit leads to a tremendous reduction in computational demand since the matrix ***H*** in Eq 7 only needs to be computed once during the Bayesian calibration stage. After initially computing ***H***, simulated RBC velocities, for a given configuration of boundary pressures, can be expediently computed by first evaluating Eq 7 and then scaling the resulting blood flows to obtain velocities as described in Section 1 in S1 File.

A well-established phase separation model was used to model the biphasic nature of blood. This model has been rigorously established based on comprehensive measurements of velocity and hematocrit in mesenteric networks[44], further refined for network simulations[30, 39, 41], and proven its applicability in a diverse set of networks ranging from artificial networks[25], fabricated microchannel networks[59, 60], mesenteric networks[28, 41], to extensive cortical networks[26, 29, 30, 36]. The convergence of the procedure for hematocrit iterations was carefully assessed, and the approach proved good convergence performance when exposed to pressure boundary conditions inferred by our Bayesian calibration framework (Table 15 in S2 file and Fig 8 to 11 in S3 File). Furthermore, relaxing the convergence criteria and allowing for a few (ten) segments to exceed the convergence threshold, led to a substantial reduction in the number of iterations needed to reach convergence without notably influencing resulting blood flow and hematocrit predictions, Table 15 in S2 File and Fig 8 to 12 in S3 File. Although the results presented are based on the most stringent convergence threshold, with no super-threshold segments allowed, future studies may benefit from the result demonstrating that mildly relaxing the convergence criteria can accelerate computations substantially without sacrificing accuracy. Incorporating the phase separation model into blood flow simulations resulted in considerable heterogeneity in hematocrit throughout the microvascular networks (Fig 8 and Fig 15 to 16 in S3 file), in agreement with previously reported results in cortical networks[26, 29, 30].

The adoption of the two-step approach for hematocrit modeling in the present study is supported by our previous observation in mesenteric networks with extensive measurements of both velocity and hematocrit available and where the phase separation effect was included into the Bayesian calibration stage[26]. However, inlet hematocrit only exhibited widespread systematic deviations from reference levels (prior means) when hematocrit measurements were included in model calibration. Since the currently studied networks lacks hematocrit measurements, and since the study focused on pressure boundary conditions and their impact on blood flow uncertainty, the increased computational effort required when merging the two steps was judged to yield marginal gains given the scope of the study.

Three viscosity model variants were considered in the evaluation of the proposed adaptive method in combination with our Bayesian calibration framework (cal.Vitro.ABC, cal.Esl.ABC, and cal.Vivo.ABC models). This decision was made to facilitate a systematic assessment of the adaptive method for pressure boundary conditions across viscosity model variants currently applied in blood flow simulations in extensive microvascular networks[26, 29–31, 34, 36, 49, 52, 61]. Simulation models incorporating the three model variants were calibrated against the same target velocities and flow directions. It was therefore expected that predicted pressures would differ across the three models, with the lowest viscosity in the Vitro model, the highest viscosity in the Vivo model, and the Esl model falling in between[41]. The analysis results confirmed these expectations (Fig 8, Table 3 to 14 in S2 File, and Fig 15 to 16 in S3 File). Shear stress, proportional to the pressure difference across a vessel segment, showed a similar effect. Despite these systematic pressure differences, a strong correlation between segment pressure was observed across the three models (Fig 8 and Fig 15 to 16 in S3 File), demonstrating that the pressure hierarchy within the microvascular networks was preserved across the three model variants. The following two factors may contribute to this similarity: Blood flow resistance is proportional to the product of segment length and effective viscosity divided by the fourth power of segment diameter. Segment diameter thereby has a strong influence on resistance independent of viscosity[41]. Additionally, microvascular networks have a highly interconnected architecture which, together with the strong heterogeneity in segment morphology, shapes the heterogeneities of hemodynamic variables in the microcirculation[2, 5].

The consistencies, across models, in individual segment hemodynamic variables, in depth dependencies in pressure profiles, and in capillary flow patterns resulted from analyses where the proposed adaptive method for pressure boundary conditions was compared against simulations based on either reference pressure boundary conditions (Experiment 1) or where identical target RBC velocities and target flow directions were used in the Bayesian calibration across models (Experiment 2). The analysis thereby demonstrated that the proposed adaptive approach, and its incorporation into our Bayesian calibration framework, yield flow simulations that are consistent with another recent method for establishing pressure boundary conditions[31]. While these results support the validity of the proposed approach for assigning boundary conditions in the examined networks, it remains an open question whether similar consistency across viscosity formulations is achieved by other methods for assigning boundary conditions. Furthermore, a formal statistical model selection between the three viscosity models is outside the scope of this study and is challenged by the lack of hemodynamic measurements in the examined networks. A comprehensive discussion of viscosity and flow resistance in the microcirculation, including factors influencing effective viscosity, is available in the cited literature[1, 41, 42, 62].

### 4.3. Uncertainty quantification

Blood flow simulation models are influenced by various sources of uncertainties and errors[26, 28]. These include model structural errors arising from the mathematical representation of complex physiological processes, uncertainties and errors related to model parameters (for example, parameters of the phase separation and viscosity models), uncertainties and errors associated with model inputs (including boundary conditions), and measurement errors in data assimilation contexts. The present study focused on the quantitative evaluation of the proposed adaptive method for pressure boundary conditions and its integration into our Bayesian calibration framework. Consequently, the uncertainty analysis correspondingly focused on how uncertainties related to pressure boundary conditions translate into uncertainties in blood flow simulations.

A simple approach to quantify the influence of uncertainties in imposed pressure boundary conditions in forward or in-silico modeling was adopted, Section 2.4.2, and evaluated, Section 3.1.2. The advantage of this method’s simplicity is that all variables required to perform the uncertainty analysis are already estimated and thereby readily available in current blood flow simulation models in terms of the matrix ***H***, Eq 7. The only additional information needed is to define the strength of uncertainty governing individual boundary conditions. The influence of this uncertainty on, for example, blood flow uncertainty, depends on the strength of the assumed uncertainty. For example, Experiment 1 considered identical strength of uncertainty across the relative pressure boundary conditions. As a result, the standard deviation and the coefficient of variance of an individual segment’s blood flow are both proportional to the assumed standard deviation governing the boundary pressures. More advanced parameterizations of boundary condition uncertainty could be implemented, such as spatially varying uncertainty profiles across boundary nodes. However, a comprehensive evaluation of such an approach is beyond the scope of this study. Despite its simplicity, the probabilistic approach to uncertainty analysis in forward models, Section 2.4.2, proved to be an effective and useful approach for quantifying the impact of pressure boundary condition uncertainty on model predictions. The approach is useful as it provides information on the reliability of hemodynamic variables and offers means for complementing hemodynamic predictions with uncertainty estimates. These uncertainty estimates could, for example, be utilized as an objective criterion for selecting segments for further analysis, thereby reducing or eliminating the impact of vessels strongly influenced by boundary condition uncertainty on summary statistics.

In Bayesian calibration, uncertainty is naturally incorporated in terms of prior distributions governing unknown parameters. By assimilating model predictions with observations, Bayes’ rule provides means for translating these uncertainties into a probability distribution governing the parameters, conditioned on the observations. Observations, of for example blood flow directions and velocities, thereby offer means for reducing the uncertainty by integrating observed behavior of the system. In this study, diameter-dependent target RBC velocities and anticipated flow directions were utilized in a subset of segments due to the lack of actual measurements in the examined networks. The aim was thereby to calibrate the models against characteristic data, given the currently available information from the literature and the reference data set, rather than calibrating the models to exactly match segment-wise hemodynamics of the reference model simulations. Despite this, strong consistencies between the reference models and the calibration models in hemodynamic estimates, as well as in depth-dependent pressure drop and layer-wise capillary flow phenomena, were observed, Fig 7 and 9. A strong agreement between layer-wise blood flow uncertainty profiles was similarly observed, Fig 2 and 6. Our Bayesian calibration framework, with the proposed adaptive method for pressure boundary conditions incorporated into it, thereby proved its validity and scalability for blood flow simulation and uncertainty quantification in high-dimensional settings with thousands of segments and unknown boundary conditions.

Although the present study focused on the impact of boundary condition uncertainty on blood flow uncertainties, it should be noted that uncertainties related to other hemodynamic variables, such as blood flow velocities, node pressures and segment pressures, and shear stress, are also readily obtainable. In linear forward modeling, estimates of uncertainty for these variables can be derived by following the approach in Eqs 7 to 11 for the respective variables. In Bayesian calibration utilizing MCMC sampling, uncertainty estimates related to such hemodynamic variables are readily available in terms of their variability across model simulation based on the MCMC samples.

Although unknown boundary conditions can be imposed by interpolating literature-derived data, by numerical optimization techniques, or by embedding or connecting techniques, they remain uncertain. For example, literature-derived data or observed data used in model fitting is likely to be corrupted by some amount of physiological variability and measurement error, which consequently impacts the specific fit. Embedding and connecting techniques are likely to depend on the actual truncation of the microvascular network at hand as well as by the specific spatial location of the network’s embedding within the larger synthetic networks or on the specific connectivity of vessels across domain boundaries. In these applications, uncertainty analysis may also be a useful approach, enabling a quantitative assessment of the reliability of hemodynamic predictions and guiding an objective selection of part of the simulated hemodynamic variables for further analysis.

### 4.4. Limitations and future research

The primary limitation of this study is the lack of measured hemodynamic data, such as blood flow rates or RBC velocities, in the examined cortical networks. Instead, linear fits to literature-derived data from awake imaging in mice[54] and RBC velocities from the reference data set[31] were used to define diameter-dependent target velocities in Experiment 2. While these data sources provide a reasonable basis for the quantitative evaluation of the proposed adaptive method for pressure boundary conditions and its integration into our Bayesian calibration framework, future extensive measurements of hemodynamic variables in extensive microvascular networks will support the continued development of the methodology.

Another limitation to the study is the uncertainty related to vessel diameters and vessel type categorization as previously discussed[31]. The rheological descriptions and the resulting hydraulic resistances strongly depend on vessel diameter, especially for small diameters, and hence significantly influence hemodynamic simulations[28, 29, 42, 62]. Vessel tortuosity was considered in calculating segment lengths, whereas uniform diameters along individual segments were used[31]. These capillary diameters have previously been adjusted to match empirical distributions while maintaining the hierarchy of vessel diameters[31]. Summary statistics of hemodynamic predictions in the reference data set have previously been evaluated against current experimental data[31]. Since the comprehensive quantitative evaluation of hemodynamic simulations against the reference data in the present study demonstrated strong correspondence between these, it is deduced that the hemodynamic predictions presented in the present study agrees with currently available data from experimental and simulation studies. Future data sets with extensive measurements, including those in smaller vessels, will enable a more comprehensive segment-wise quantitative model evaluation of hemodynamic predictions against actual measurements, as previously done in other tissues[26, 28, 39, 41]. Additionally, such data could further facilitate the incorporation of uncertainties in vessel diameters, for example, into the calibration analysis and the adaptation of these diameters to align model predictions with measurements[27].

Whereas this study primarily focused on the influence of boundary pressure uncertainty on model simulations, future studies could expand to include other sources of uncertainties, such as those related to diameters, as discussed above, or to the parameters of the rheological descriptions including the phase separation model[39]. Additionally, future analyses could also further investigate the effects of measurement uncertainty and consider more advanced statistical error models[26]. Again, these future research directions will be supported by future comprehensive measurements of hemodynamic variables in extensive networks.

In summary, a methodical approach, with increasing model complexity, was employed to establish a link between this study and previously published results[31]. An adaptive method for selecting appropriate pressure boundary conditions was proposed and rigorously tested against predictions of an established blood flow simulation model[25, 31]. The adaptive method was further integrated into our Bayesian calibration framework[26], which provided hemodynamic simulations in strong agreement with reference model simulations. Importantly, the present study demonstrated that previously published results of depth-dependent pressure profiles and layer-wise capillary RBC flow patterns[31], obtained with an in-vitro viscosity formulation, generalize for in-vivo viscosity formulations and across the examined approaches for selecting pressure boundary conditions. The impact of pressure boundary condition uncertainty on blood flow uncertainty was quantitatively evaluated using a probabilistic approach in forward modeling and Bayesian calibration in inverse modeling. The uncertainty quantification revealed a spatially varying pattern in blood flow uncertainty. In conclusion, it is anticipated that the adaptive method for pressure boundary conditions will be useful in future applications of both forward and inverse blood flow modeling, and that uncertainty quantification will be valuable in complementing hemodynamic predictions with associated uncertainties.

## 5. Appendix

### 5.1. Relationship with sensitivity analysis

The partial derivatives of blood flows ***q*** in Eq (8) with respect to the uncertain boundary variables in

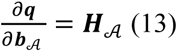

Consequently, the individual columns in ***H***_𝒜_ quantify the sensitivity of segment blood flows with respect to the corresponding boundary variables (in columns). In case of a diagonal covariance matrix ***Σ***_𝒜_, it follows that the diagonal of the covariance matrix ***Ω*** in Eq 11 is equivalent to first squaring the sensitivities in Eq (13), then scaling (multiplying) each column according to the variances found along corresponding diagonal element of ***Σ***_𝒜_, and finally summing across columns to accumulate contributions of individual scaled sensitivities. The squaring of individual sensitivities is commonly done in sensitivity analysis to avoid cancellation effects when accumulating across single sensitivities. The partial derivative in Eq (13) quantifies changes in blood flow per unit change in the boundary variables. Scaling by the variances associated with individual boundary variables is therefore useful since it allows assigning varying levels of uncertainty across the uncertain boundary variables. Furthermore, it allows boundary variables with different physical units to be considered, which would be the case if both pressure boundary conditions and flow boundary conditions are modeled as uncertain parameters.

### 5.2. Software and data availability

The methods for blood flow simulation, uncertainty quantification, and Bayesian calibration described in this paper are implemented in C++ and will be available in a publicly available open-source software suite, NetInf, at GitHub <GitHub URL placeholder, will be available upon acceptance of this manuscript>. The repository contains source code, build instructions, vignettes for analyses, and example data. Data files with blood flow simulation results, underlying the presented summary figures and tables, and analysis scripts for reproducing figures and tables from these data files will be available <Zenodo URL placeholder, will be deposited upon acceptance of this manuscript>.

## 6. Supporting information

**S1 File. Supplementary text.** (PDF)

**S2 File. Supplementary tables.** (PDF)

**S3 File. Supplementary figures.** (PDF)

## 7. Acknowledgements

The author would like to thank Franca Schmid and her colleagues for making the microvascular networks and corresponding model simulations publicly available.

